# Mitochondrial Ca^2+^ influences photoresponse recovery, metabolism, and mitochondrial localization in cone photoreceptors

**DOI:** 10.1101/510792

**Authors:** Rachel A. Hutto, Celia M. Bisbach, Fatima Abbas, Daniel Brock, Whitney M. Cleghorn, Edward D. Parker, Benjamin Bauer, William Ge, Frans Vinberg, James B. Hurley, Susan E. Brockerhoff

## Abstract

Mitochondrial Ca^2+^ regulates key cellular processes including cytosolic Ca^2+^ signals, energy production, and susceptibility to apoptosis. Photoreceptors are specialized neurons that have extraordinarily high energy demands and rely on cytosolic Ca^2+^ signals for light adaptation and neurotransmission. Here we show that unlike other neurons zebrafish cone photoreceptors express low levels of the mitochondrial Ca^2+^ uniporter (MCU), the channel that allows mitochondrial Ca^2+^ entry. To determine why MCU expression is kept low, we overexpressed MCU specifically in cones. This increases mitochondrial [Ca^2+^], causes faster cytosolic Ca^2+^ clearance, and accelerates photoresponse recovery. Moreover, flux through the citric acid cycle increases despite dramatic changes in mitochondrial ultrastructure and localization. Remarkably, cones survive this ongoing stress until late adulthood. Our findings demonstrate the importance of tuning mitochondrial Ca^2+^ influx to modulate physiological and metabolic processes and reveal a novel directed movement of abnormal mitochondria in photoreceptors.

## Introduction

Photoreceptors are highly specialized sensory neurons responsible for vision. In addition to having unique and vulnerable structural features, photoreceptors reside in the retina, a hostile cellular environment. They can be exposed to damaging light radiation, are located near blood vessels with high levels of oxygen, and use more ATP than most cells in the body. Despite these chronic stressors, most people retain vision throughout their lives, highlighting the extraordinary ability of photoreceptors to regulate cellular homeostasis and maintain viability.

Photoreceptor function and survival depends on Ca^2+^ homeostasis. Photoreceptors rely on Ca^2+^ as a second messenger to recover from light signals and adapt to constant illumination (Nakatani and Yau, 1988). In darkness they continuously release synaptic vesicles, which requires precise regulation of synaptic Ca^2+^ by L-type voltage-gated channels (Barnes and Kelly, 2002). Both chronic elevations and chronic decreases in cytosolic Ca^2+^ have been implicated in photoreceptor cell death and retinal disease (for review, see Fain, 2006; Vinberg et al., 2018). Mutations in photoreceptor phosphodiesterase, guanylyl cyclase, and guanylyl cyclase activating protein result in sustained high Ca^2+^ in the cell and cause retinal degeneration (Bowes et al., 1990; Dizhoor et al., 1998; Fox et al., 1999; Payne et al., 1998; Sokal et al., 1998; Tucker et al., 1999; Wilkie et al., 2000). Sustained light exposure or deficiencies of rhodopsin kinase and arrestin that cause sustained low intracellular Ca^2+^ also cause degeneration of photoreceptors (Chen et al., 1999a, 1999b; LaVail et al., 1987).

Cytosolic Ca^2+^ in photoreceptors is buffered and regulated by the endoplasmic reticulum (ER) and mitochondria (for review, see Križaj, 2012). In the synapse, Ca^2+^ flow through the ER to the synaptic terminal supports sustained CICR-driven synaptic transmission (Chen et al., 2015). Mitochondria are most abundant in a region of the cell body termed the “ellipsoid”, between the outer segment (where phototransduction occurs) and the rest of the cell body and synapse (Hoang et al., 2002). Ca^2+^ pools in the outer segment are distinct from the rest of the cell, and it has been suggested that ellipsoid mitochondria can protect the cell body from the high Ca^2+^ generated in the outer segment as a consequence of phototransduction (Križaj and Copenhagen, 1998; Szikra and Krizaj, 2007). It was recently shown in zebrafish photoreceptors that ellipsoid mitochondria maintain the distinct Ca^2+^ pools between the outer segment and the cell body (Giarmarco et al., 2017).

Mitochondrial Ca^2+^ uptake is also involved in energetic output (Glancy and Balaban, 2012). Ca^2+^ can regulate the activity of mitochondrial enzymes including pyruvate dehydrogenase, isocitrate dehydrogenase, and α-ketoglutarate dehydrogenase (Denton, 2009). The aspartate/glutamate exchangers on the inner mitochondrial membrane are sensitive to Ca^2+^ (Contreras et al., 2007; Satrústegui et al., 2007). Ca^2+^ also may regulate the activity of ATP synthase (for review, see Glancy and Balaban, 2012). Increased mitochondrial matrix Ca^2+^ has been linked to increases in NADH and ATP production in multiple cell types (Balaban, 2009; Hajnóczky et al., 1995; Jouaville et al., 1999). However, metabolic responses to changes in mitochondrial Ca^2+^ vary across tissues, likely reflecting the diverse metabolic demands of different tissues (Griffiths and Rutter, 2009). It is not currently known to what extent fluxes in mitochondrial Ca^2+^ influence metabolism in photoreceptors, which have extraordinarily high ATP demand (Okawa et al., 2008).

While modest increases in mitochondrial Ca^2+^ can stimulate energy production, excessive mitochondrial Ca^2+^ can trigger opening of the mitochondrial permeability transition pore (Baumgartner et al., 2009). Prolonged opening of the pore is associated with mitochondrial swelling, collapse of the proton gradient, and release of mitochondrial solutes that eventually results in cell death (Bernardi et al., 1999; Kroemer et al., 1997; Rizzuto et al., 2012; Smaili et al., 2000; Zoratti and Szabò, 1995). In isolated rat retina, increases in cellular Ca^2+^ and Pb^2+^ cause photoreceptor-selective apoptosis that depends on mitochondrial permeability transition pore activity (He et al., 2000).

Ca^2+^ import into the mitochondrial matrix in all cells occurs via the mitochondrial Ca^2+^ uniporter complex (MCU complex), comprised of a tetramer or pentamer of the pore-forming protein MCU that associates with several regulator proteins (Baradaran et al., 2018; Baughman et al., 2011; Fan et al., 2018; Nguyen et al., 2018; Oxenoid et al., 2016; De Stefani et al., 2011, 2015; Yoo et al., 2018). The small transmembrane protein EMRE is necessary for MCU function in vertebrates (Sancak et al., 2013). MICU proteins tune Ca^2+^ uptake through the uniporter complex. MICU1 imparts cooperativity to Ca^2+^ entry and is required for other MICU proteins to interact with the MCU complex (Csordás et al., 2013; Kamer et al., 2014; Mallilankaraman et al., 2012). MICU2 has a distinct inhibitory effect on Ca^2+^ entry at high cytosolic Ca^2+^ concentrations (Patron et al., 2014; Plovanich et al., 2013). MICU3, enriched in neurons, confers hyper-Ca^2+^ uptake activity (Patron et al., 2018). The MCU pore can also include a dominant negative form of the MCU subunit, called MCUb (Raffaello et al., 2013). Finally, several other mitochondrial proteins have been purported to interact with the MCU complex and potentially modulate its function (Chaudhuri et al., 2016; Hoffman et al., 2014; Nieminen et al., 2014; Paupe et al., 2015; Tomar et al., 2016; Zeng et al., 2018). This degree of regulation of MCU, along with the variability of modulator expression across tissues, implies that the activity of this complex is attuned specifically to cellular needs.

To begin to investigate the relationship between Ca^2+^ and photoreceptor mitochondria we analyzed protein and transcript levels of MCU, MICU1, MICU2 and MICU3 in retina compared to heart and brain. We found that MICU transcript expression in retina and brain is strikingly similar. However, the retina has much lower MCU protein expression than brain or heart tissue, and MCU is particularly low in cone photoreceptors. To elucidate why MCU expression may be limited in cones, we generated zebrafish models of cone-specific MCU overexpression. We found that elevated MCU in cones increases mitochondrial Ca^2+^ content, alters cytosolic Ca^2+^ transients and the photoresponse, and increases flux through citric acid cycle enzymes. MCU overexpression also causes dramatic changes in mitochondrial morphology accompanied by a directed movement of mitochondria away from the ellipsoid region of the cell. Despite these morphological changes and disruptions to Ca^2+^ and cellular homeostasis, cones overexpressing MCU survive well into adulthood before degenerating. Our findings uncover how mitochondrial Ca^2+^ content can affect cone function and open a new area of research investigating survival strategies employed by these sensory neurons.

## Results

### MCU expression is specifically limited in retinal tissue

We developed a custom antibody against the purified N-terminus of zebrafish MCU and verified the specificity of this antibody using a global zebrafish MCU knock-out (MCU KO, **Figure 1A**). Using this antibody, we found that MCU protein expression (as normalized to total protein) is significantly higher in brain tissue than retina, which is even lower than heart tissue (**Figure 1A**). When normalizing MCU protein to other mitochondrial inner membrane proteins cytochrome oxidase (MTCO1) and succinate dehydrogenase (SDH), the retina more closely resembles heart tissue (**Figure 1B**). To see if this low retinal abundance of MCU was reflected in other regulators of the complex, we used RT-qPCR to determine the relative mRNA quantity of the MCU regulators MICU1, MICU2, and MICU3 (which has two isoforms in zebrafish) across the same tissues. Expression of all MICU transcripts in retina is more similar to expression in brain rather than heart (**Figure 1C**).

**Figure 1:**
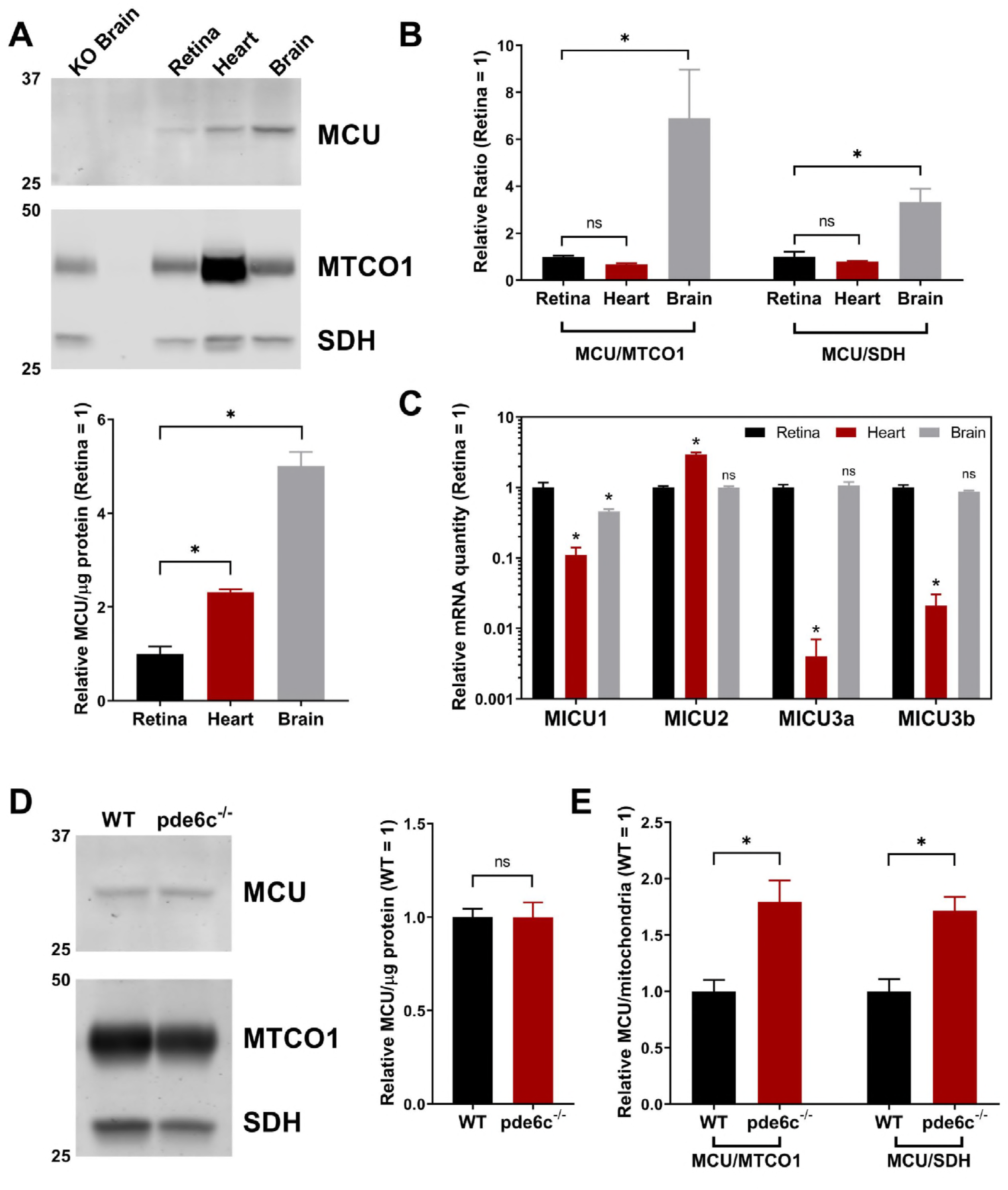
MCU expression is limited in the retina, particularly in cones. A. Western blot using zebrafish MCU antibody with zebrafish tissue lysate, enriched for mitochondrial proteins and probed for MCU, mitochondrial cytochrome oxidase (MTCO1), and succinate dehydrogenase (SDH). Samples were pooled from either 4 retinas, 2 hearts, or 1 brain and performed with n=3 biological replicates of each pool. Each lane contains 8 μg of protein. Values were normalized relative to retina tissue. Bars = standard error. *p<0.05 using ANOVA followed by Dunnett post-hoc test (comparison to retina). B. Within each lane from the gel in A, the ratios of MCU signal to the mitochondrial proteins MTCO1 and SDH were determined. Values were normalized relative to retina tissue. Bars = standard error. *p<0.05 using ANOVA followed by Dunnett post-hoc test (comparison to retina). C. qRT-PCR quantification of relative mRNA of MICU proteins (relative to reference gene *Ef1α* and/or *b2m*, see methods) across retina, heart, and brain tissues. Bars = standard error. *p<0.05 and ns = not significant using ANOVA followed by Dunnett post-hoc test (comparison to retina). D. Retinal lysate, enriched for mitochondrial proteins, of WT and pde6c^−/−^ cone deficient retinas. Each lane is from mitochondrial-enriched lysate of two pooled retinas from a single fish, n=4 fish. Each lane contains 30 μg of protein. Ns = not significant using Welch’s t-test. E. Relative quantification of the ratio MCU to mitochondrial proteins MTCO1 and SDH in WT and pde6c^−/−^ cone deficient retinas from the gel shown in D. *p<0.05 using Welch’s t-test.

To quantify MCU expression specifically in zebrafish cone photoreceptors, we took advantage of a zebrafish cone degeneration model caused by a mutation in the cone phosphodiesterase (pde6c^−/−^); in this model, cones selectively degenerate while rods and other retinal neurons are preserved (Stearns et al., 2007). We analyzed MCU and mitochondrial membrane protein expression in the pde6c^−/−^ mutants compared to their WT siblings and found that without cones there is a loss in mitochondrial membrane protein expression that is not reflected in a loss of MCU signal (**Figure 1D,E**). The loss of MTCO1 and SDH expression in the pde6c^−/−^ mutant is consistent with cones having more mitochondrial volume than rods (Hoang et al., 2002), but the data indicate that these cone mitochondria have relatively little MCU compared to other retinal neurons. Our antibody was not suitable for IHC of the endogenous protein.

### Overexpression of MCU in cones raises basal [Ca^2+^] in the mitochondrial matrix

We hypothesized that low expression of MCU protein in photoreceptors could protect them from overloading their mitochondria matrix with Ca^2+^. To test the consequences of increased MCU, we expressed MCU-T2A-RFP in zebrafish cones under control of the promoter for cone transducin (“*gnat2*” or “TαCP”) and established a stable transgenic line overexpressing zebrafish MCU in cone mitochondria (**Figure 2A**). Cones overexpressing MCU (MCU OE) also express RFP in the cytosol, as the T2A sequence stalls the ribosome after MCU has been translated to allow MCU protein release before translation of RFP (Kim et al., 2011). Based on immunoblot analysis we found that retinal MCU protein expression is 102 ± 5-fold higher in the MCU OE model. This also is reflected in the ratio of MCU to other mitochondrial membrane proteins (**Figure 2B,C**). Immunohistochemistry confirmed that MCU expressed in this way is localized to cone mitochondria (**Figure 2D**).

**Figure 2:**
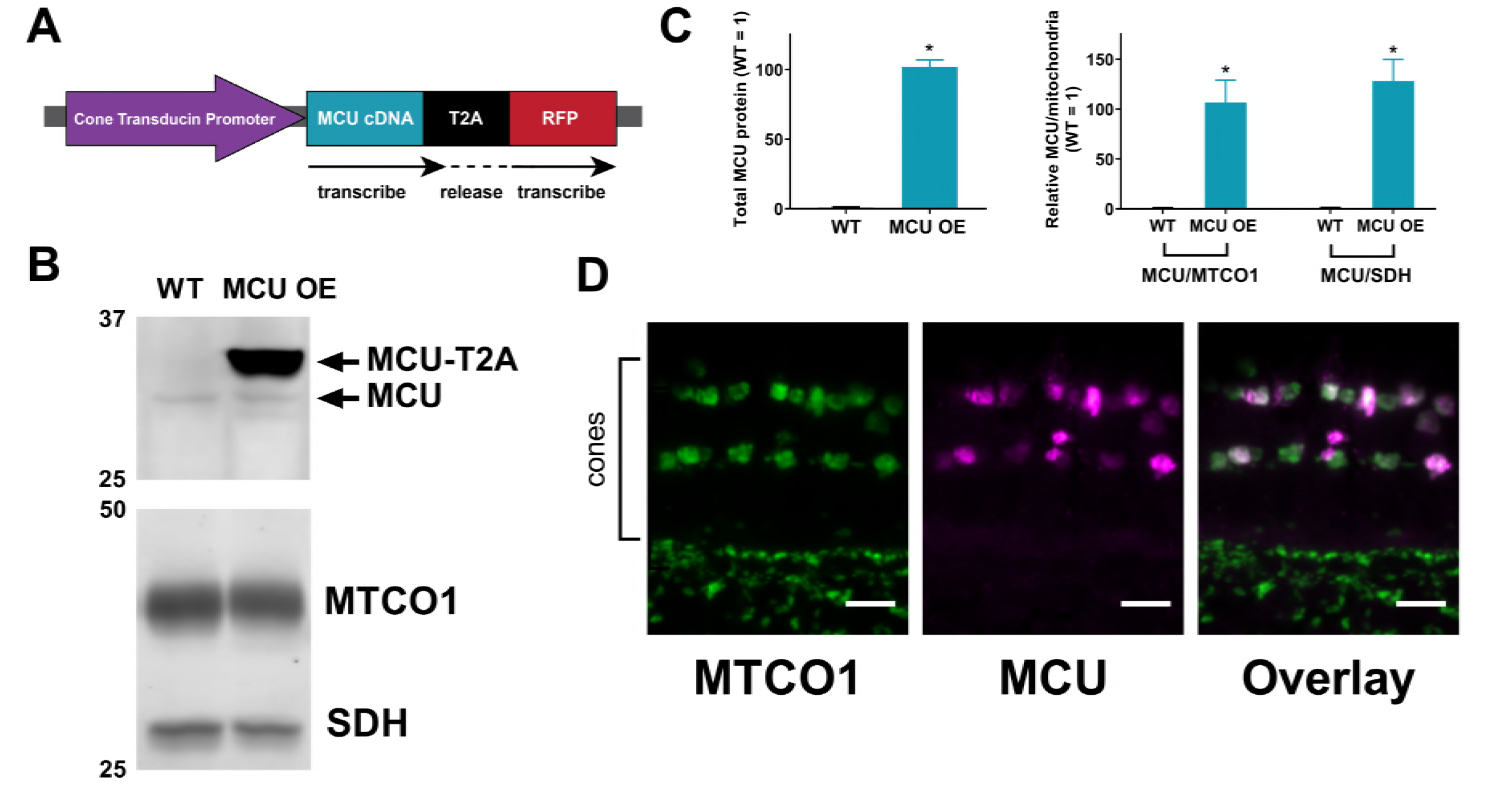
Successful generation of a cone-specific MCU overexpression zebrafish model. A. Schematic of MCU OE construct. The cone transducin promoter (TαCP, *gnat2*) drives expression of zebrafish MCU cDNA in all cone subtypes. The MCU cDNA is tagged with a T2A sequence followed by RFP. The T2A sequence causes ribosomes to stall and release the nascent MCU polypeptide with some added peptides from the T2A sequence before translating the RFP separately. Thus, RFP is present in the cytosol of cones with MCU overexpression. B. Retinal lysate, enriched for mitochondrial proteins, of WT and MCU OE retinas. Blot was probed with antibodies for MCU, MTCO1, and SDH. 8 μg of protein was loaded per well. Each lane is from mitochondrial-enriched lysate of two pooled retinas from a single fish, n=4 fish. C. Quantification of relative MCU signal as a function of protein concentration and relative to other mitochondrial markers from the gel in B. Both exogenous and endogenous MCU were used for total MCU quantification in the MCU OE retina. *p<0.05 using Welch’s t-test. D. Immunohistochemistry of a larval zebrafish retina expressing the MCU construct in A using MCU and mitochondrial cytochrome oxidase (MTCO1) antibodies. Scale bar = 5 μm.

To test the effects of MCU protein overexpression on mitochondrial Ca^2+^ content, we crossed MCU OE fish with the *gnat2*:mito-GCaMP3 transgenic line, which expresses the Ca^2+^ sensor GCaMP3 in cone mitochondria (Giarmarco et al., 2017). Mito-GCaMP3 fluorescence in live zebrafish larvae is 4.4 fold higher (median, Q1: 3.4, Q3: 6.06 fold) in MCU OE models compared to WT siblings (**Figure 3A**). We next prepared *ex vivo* retinal slices (Giarmarco et al., 2018) of MCU OE gnat2:mito-GCaMP3 zebrafish and sibling controls. We measured the baseline mito-GCaMP3 fluorescence (F_0_), the maximum fluorescence (F_max_) with ionomycin and 2 mM Ca^2+^, and the minimum fluorescence (F_min_) in 5 mM EGTA (**Figure 3B**). Comparing (F_0_ − F_min_) to (F_max_ − F_min_) indicated that the baseline GCaMP signal is operating at 20 ± 1% of maximum fluorescence in WT mitochondria and 48 ± 2% of maximum in MCU OE mitochondria. Based on these measurements, we estimate that the baseline free [Ca^2+^]_mito_ is 80.0 nM (median, with Q1: 67.1, Q3: 110.5 nM) in WT mitochondria and 320.6 nM (median, with Q1: 223.9, Q3: 509.0 nM) in MCU OE mitochondria (**Figure 3C, equation in legend**).

**Figure 3:**
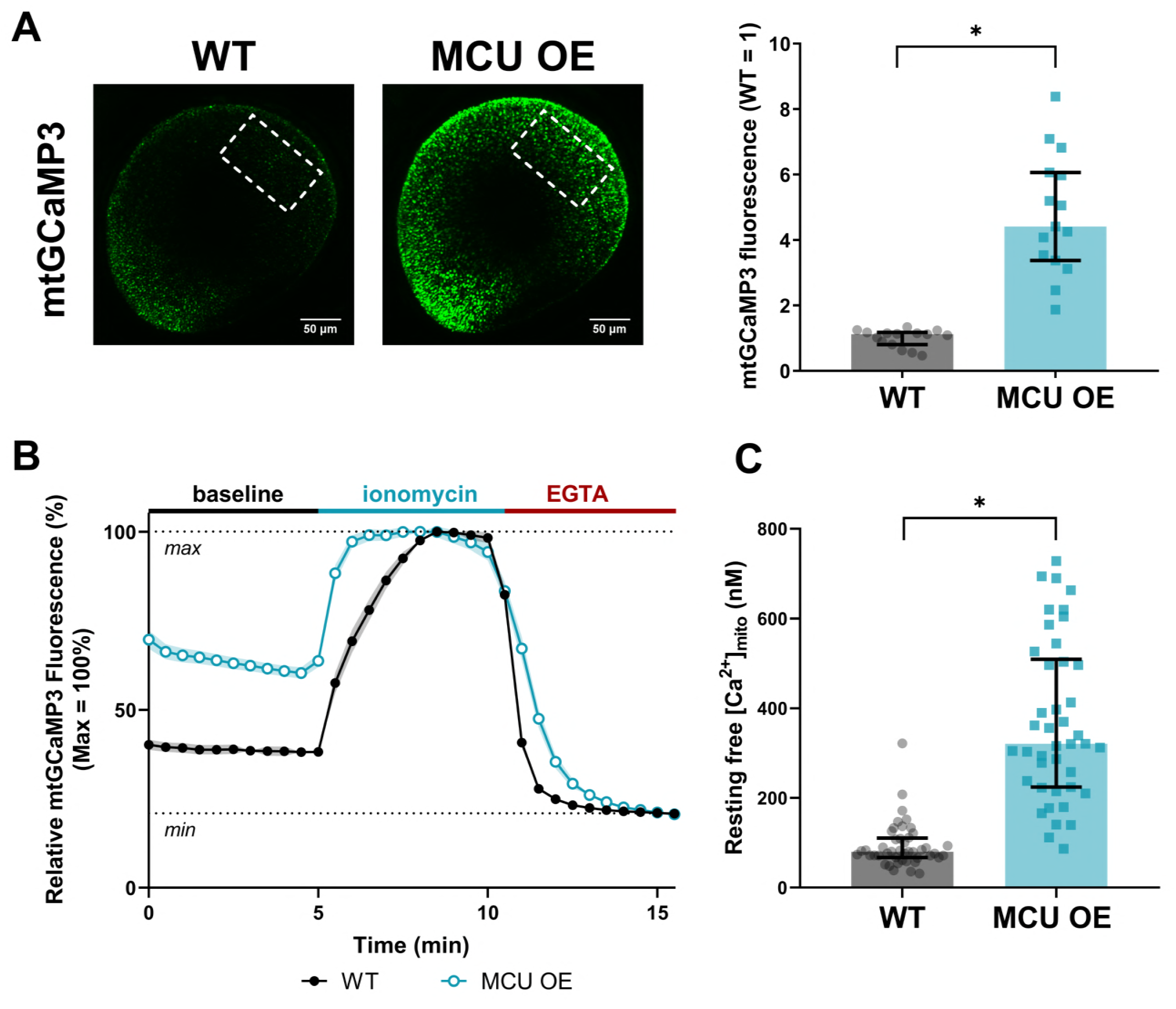
MCU overexpression in cones increases basal [Ca^2+^]_mito_. A. Total cone mitochondrial clusters in a larval zebrafish eye expressing *gnat2*:mtGCaMP3, a mitochondrial Ca^2+^ sensor (green). Dotted outlines demarcate the region of the eye used for fluorescence quantification. Reporting the median with bars = interquartile range, n=15 larvae for both WT and MCU OE. *p<0.05 using Mann-Whitney test. B. Relative mito-GCaMP3 fluorescence of cone mitochondrial clusters in adult retinal slices of *gnat2*:mtGCaMP3 fish (WT or MCU OE). Baseline fluorescence was first assayed in the presence of KRB buffer containing 2 mM CalCl_2_, then ionomycin (5 μM) was added to the slice to allow 2 mM Ca^2+^ entry into the mitochondria to saturate the probe. Next, EGTA (5 mM) was added to the solution (keeping [ionomycin] constant) to chelate Ca^2+^ and establish the minimum GCaMP3 fluorescence signal. n= 45 mitochondrial clusters (3 fish) for WT and n=42 mitochondrial clusters (3 fish) for MCU OE. Fish were between 3-5 months of age and slices were imaged every 30 s. Shaded region = standard error. C. Approximation of resting free [Ca^2+^] in mitochondrial clusters assayed in B. Approximations used the equation 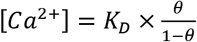, where 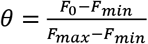. We used the previously reported K_D_ of GCaMP3 (345 nM, from Chen et al., 2013) as an approximation for our calculation. Reporting the median with bars = interquartile range and *p<0.05 using Mann-Whitney test.

### Overexpression of MCU in cones reduces cytosolic Ca^2+^ transients and alters the cone photoresponse

Increased mitochondrial Ca^2+^ uptake could alter the clearance kinetics of cytosolic Ca^2+^ transients. To test this, we prepared *ex vivo* retinal slices of control and MCU OE *gnat2*:GCaMP3 zebrafish, which express the Ca^2+^ sensor GCaMP3 in the cytosol of cones. We pre-incubated the retinal slices in a 0 mM Ca^2+^ solution before introducing a bolus of 5 mM CaCl_2_. We then used timelapse imaging to monitor the clearance of cytosolic Ca^2+^ from the cell body (**Figure 4A**). Consistent with increased Ca^2+^ uptake by mitochondria, MCU OE cones clear Ca^2+^ from the cell body cytosol 2.3 ± 0.1 times faster than their WT siblings, as determined by the decay constant of a single exponential fit (**Figure 4B, Supplemental Figure 4A**). We also found that the peak fold change in cone cell body GCaMP3 fluorescence in response to the Ca^2+^ bolus is lower in MCU OE compared to their WT siblings (**Figure 4C**). To determine whether these changes were due to Ca^2+^ uptake via MCU and not another secondary effect, we incubated MCU OE retinal slices in Ru360, a drug that blocks Ca^2+^ uptake through the MCU; these experiments significantly restored the WT phenotype (**Figure 4A-C**).

**Figure 4:**
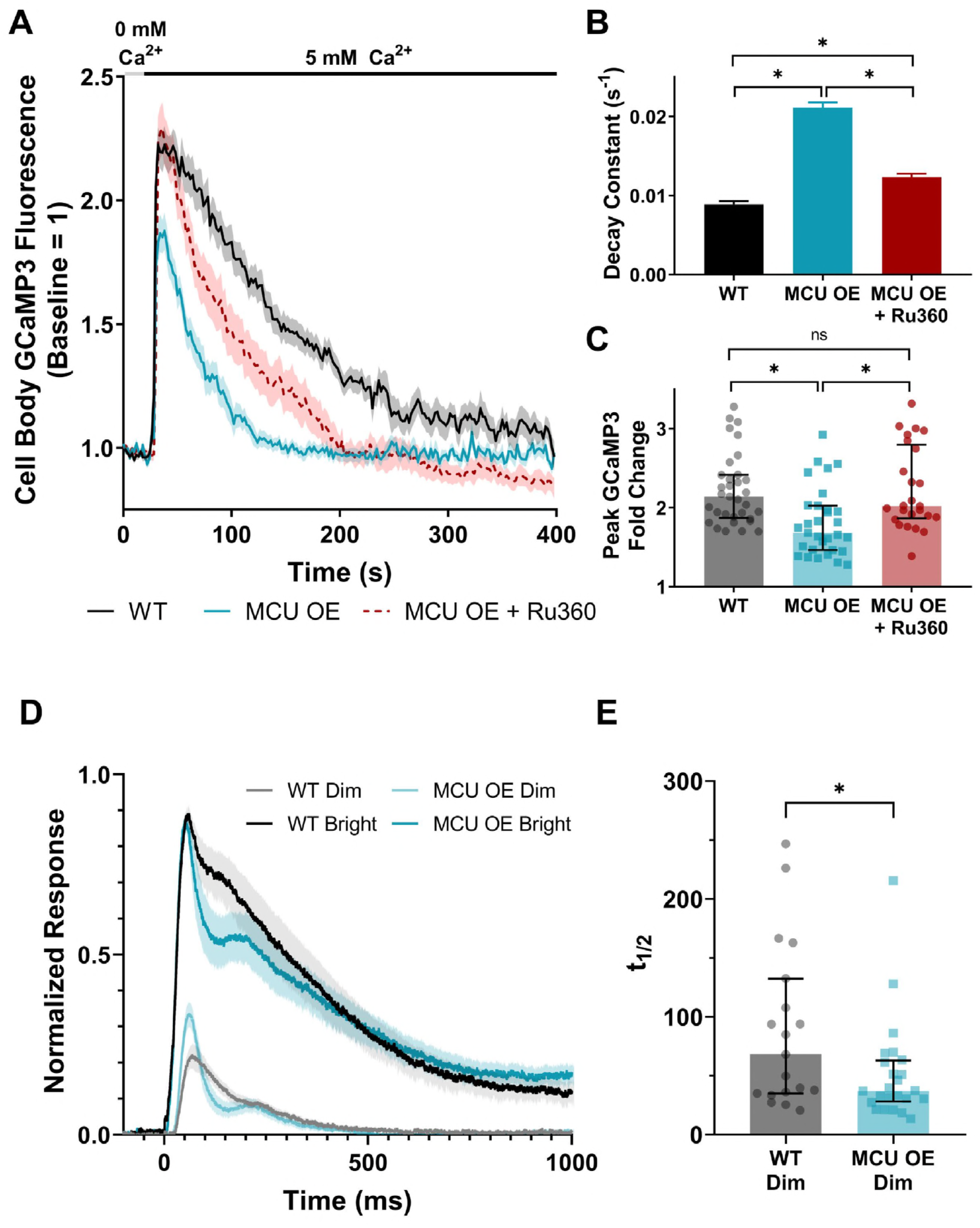
MCU overexpression in cones accelerates cytosolic Ca^2+^ clearance with functional consequences on the photoreceptor response to light. A. Isolated retinas from *gnat2*:GCaMP3 fish pre-incubated in 0 mM Ca^2+^ for 10 min then subjected to a 5 mM Ca^2+^ bolus (black bar). Fish used were WT, MCU OE, or MCU OE retinas preincubated in Ru360 (100 μM) and maintained throughout the experiment. N=33 cells (7 fish) for WT, n=31 cells (7 fish) for MCU OE, n=26 cells (3 fish) for MCU OE + Ru360. Fish were between 3-5 months of age and slices were imaged every 2s. Shaded region = standard error. B. Decay constant of Ca^2+^ clearance for experiments shown in A. Bars = standard error. *p<0.05 using ANOVA followed by Tukey post-hoc test. C. Peak GCaMP3 fluorescence fold-change from baseline for experiments shown in A. The median is reported and bars = interquartile range. WT: median=2.14, Q1=1.87, Q3=2.42. MCU OE: median=1.68, Q1=1.46, Q3=2.02. MCU OE + Ru360: median=2.02, Q1=1.87, Q3=2.80. *p<0.05 using Kruskal-Wallis followed by Dunn post-hoc test. D. The normalized *ex vivo* a-wave response isolated using DL-AP4 (40 μM) and CNQX (40 μM). Each retina response is normalized to R_max_, the maximum response at the brightest light intensity. N=19 retinas (11 fish) for WT siblings, n=24 retinas (14 fish) for MCU OE. Fish were 7 months of age. Shaded region = standard error. E. Time to half maximum of the individual responses to a dim stimulus flash from data shown in D. The median is reported and bars = interquartile range. WT: median=68.1 s, Q1=34.9, Q3=132.4. MCU OE: median=37.1 s, Q1=28.2, Q3=63.0 *p<0.05 using Mann-Whitney test.

Cytosolic Ca^2+^ in the photoreceptor outer segments regulates the gain of phototransduction, light response recovery kinetics and light adaptation mainly via GCAPs- and recoverin-mediated pathways both in rods and cones (Makino et al., 2004; Mendez et al., 2001; Sakurai et al., 2011, 2015). Compared to rod photoreceptors, cone phototransduction gain is smaller and cones recover faster to the dark-adapted state after transient light stimulus (for recent review, see Vinberg et al., 2018). These differences are believed to stem, at least partly, from the faster clearance of cytosolic Ca^2+^ in cone as compared to rod outer segments (Sampath et al., 1998, 1999). We asked whether overexpression of MCU would contribute to a faster clearance of Ca^2+^ from the cone outer segments and consequently lead to a faster photoresponse recovery. We used an ex vivo ERG technique to measure pharmacologically isolated cone photoreceptor responses from adult WT and MCU OE retinas and found that the initial phase of photoreceptor response recovery following light flashes is accelerated by MCU overexpression (**Figure 4D**). This is most apparent in the dim flash responses, which have a shorter time to half-maximum in MCU OE retinas (**Figure 4E**). At this age (7 months) the maximal response amplitude (R_max_) is somewhat decreased (**Supplemental Figure 4B**). However, the dim flash responses normalized to R_max_ increase, suggesting a higher gain of phototransduction in MCU OE cones (**Figure 4D, Supplemental Figure 4C**).

### Retinas with MCU overexpressing cones have altered flux through the TCA cycle

It has been reported that pyruvate dehydrogenase (PDH), isocitrate dehydrogenase (IDH), and α-ketoglutarate dehydrogenase (α-KGDH) have higher activity in the presence of Ca^2+^ (Denton and Randle 1972, McCormack and Denton 1979, McCormack and Denton 1990). Ca^2+^ can boost PDH activity by stimulating PDP1c, which is a subunit of the phosphatase that converts inactive phosphorylated PDH to active unphosphorylated PDH (Denton 2009). How Ca^2+^ stimulates α-KGDH and IDH is not as well understood, but it is thought that Ca^2+^ increases activity by interacting directly with these enzymes to lower the K_M_ for their substrates (Denton 2009).

We investigated the effect of MCU overexpression on PDP1c phosphatase activity by immunoblotting WT and MCU OE retinas with antibodies against phosphorylated PDH (P-Ser293 of E1α) and against total E1α. While we hypothesized MCU OE cones would have a lower P-PDH/total PDH ratio, we observed that it was instead very slightly increased (**Figure 5A, Supplemental Figure 5A**).

To investigate the effects of MCU overexpression on α-KGDH and IDH, we performed a series of metabolic flux experiments. Cones rely on glucose to fuel both aerobic glycolysis and the TCA cycle in order to meet the high energy demands required for phototransduction. We incubated WT and MCU OE retinas in U-^13^C-glucose, extracted metabolites, and used gas chromatography-mass spectrometry (GC-MS) to quantify ^13^C-labeled metabolites in these pathways (**Figure 5B**). Flux through glycolysis and total metabolite levels are largely unaltered in MCU OE cones (**Supplemental Figure 5C,D**). However, the time course of ^13^C-metabolite accumulation indicates that IDH and α-KGDH activities are enhanced (**Figure 5C**). The steady state levels of m2 citrate and m2 isocitrate are lower in MCU OE retinas compared to WT retinas. This is consistent with Ca^2+^ lowering the K_m_ of IDH, leading to depletion of upstream metabolites. High IDH activity alone should cause more m2 α-KG accumulation, but instead we observe similar steady state levels of m2 α-KG in MCU OE retinas and an accumulation of metabolites downstream of α-KGDH. This suggests that MCU OE retinas also have increased α-KGDH activity which is preventing the buildup of α-KG. Overall, these results are consistent with Ca^2+^ increasing activity of both IDH and α-KGDH.

**Figure 5:**
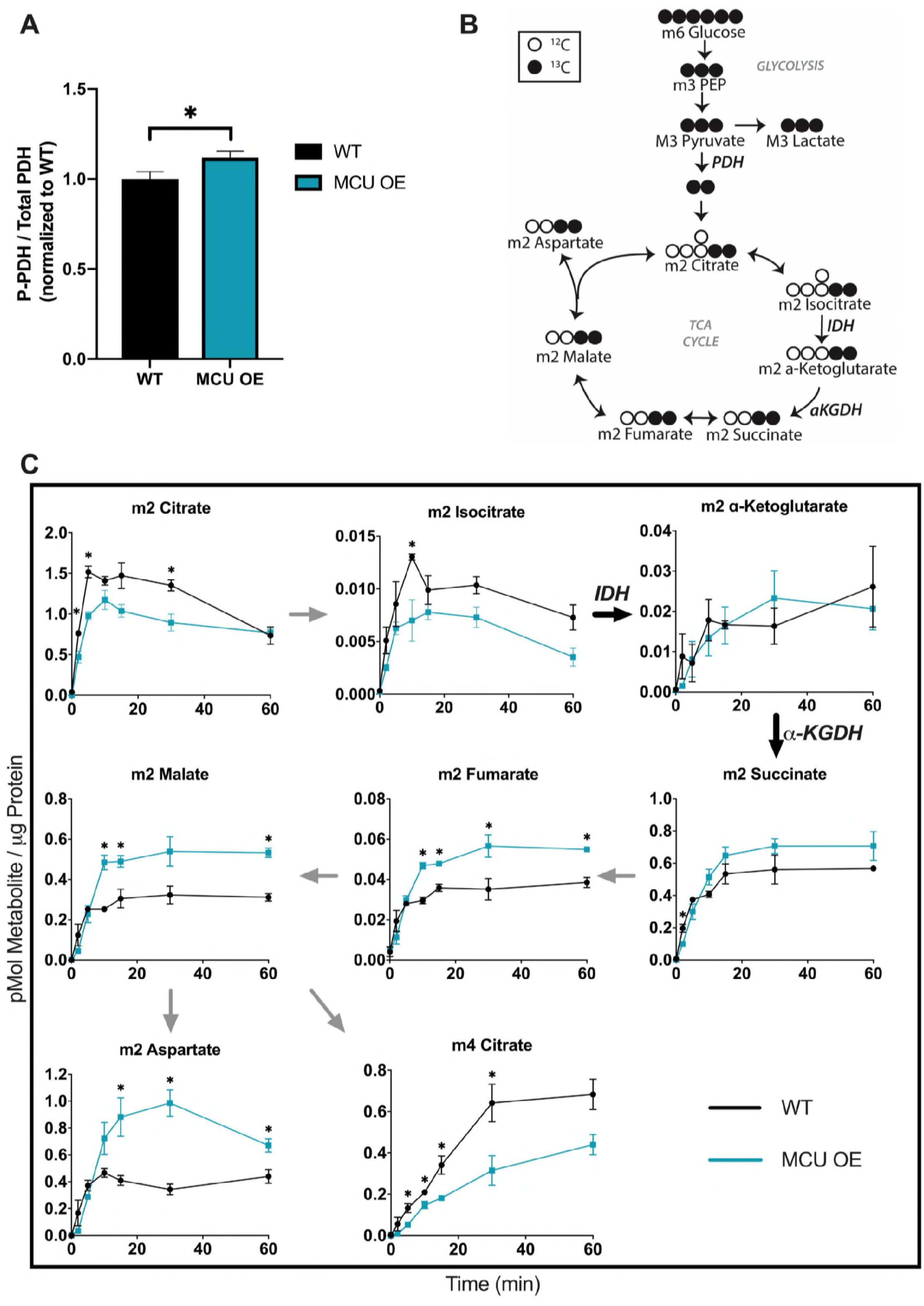
^13^C-Glucose reveals increased flux through the TCA cycle in MCU OE retinas. A. P-PDH/total PDH ratio from WT and MCU OE retinas, normalized to WT. MCU OE retinas have a 1.12 ± 0.02-fold higher P-PDH/total PDH ratio than WT. n=6 WT and 6 MCU OE retinas. *p<0.05 using Welch’s t-test. B. Diagram showing how labelled carbons from U-^13^C-glucose are incorporated through glycolysis and the first round of the TCA cycle. Shaded = labeled carbon, empty = unlabeled carbon. C. Levels of isotopomers in WT and MCU OE retinas. ‘m’ signifies the number of ^13^C-labeled carbons in each metabolite. ‘m2’ TCA cycle metabolites are made from one round of the TCA cycle. Data points represent averages from n=3 retinas from 3 different fish. *p<0.05 using Welch’s t-test.

While glucose is a physiologically relevant fuel for photoreceptors, it makes it difficult to observe IDH and α-KGDH activity in isolation because the two are intrinsically linked in the TCA cycle. For this reason, we next used U-^13^C-glutamine in the absence of glucose to bypass IDH and directly fuel α-KGDH (**Figure 6A**). Since glutamine is not a typical fuel for photoreceptors, we tested concentrations of ^13^C-glutamine ranging from 0.1 mM to 2 mM (**Supplemental Figure 6A**). We found that at most concentrations of ^13^C-Glutamine, MCU OE retinas produced metabolites downstream of α-KGDH at higher levels compared to WT (**Figure 6B**). Increased production of these metabolites further confirms that α-KGDH activity is higher in MCU OE cones.

**Figure 6:**
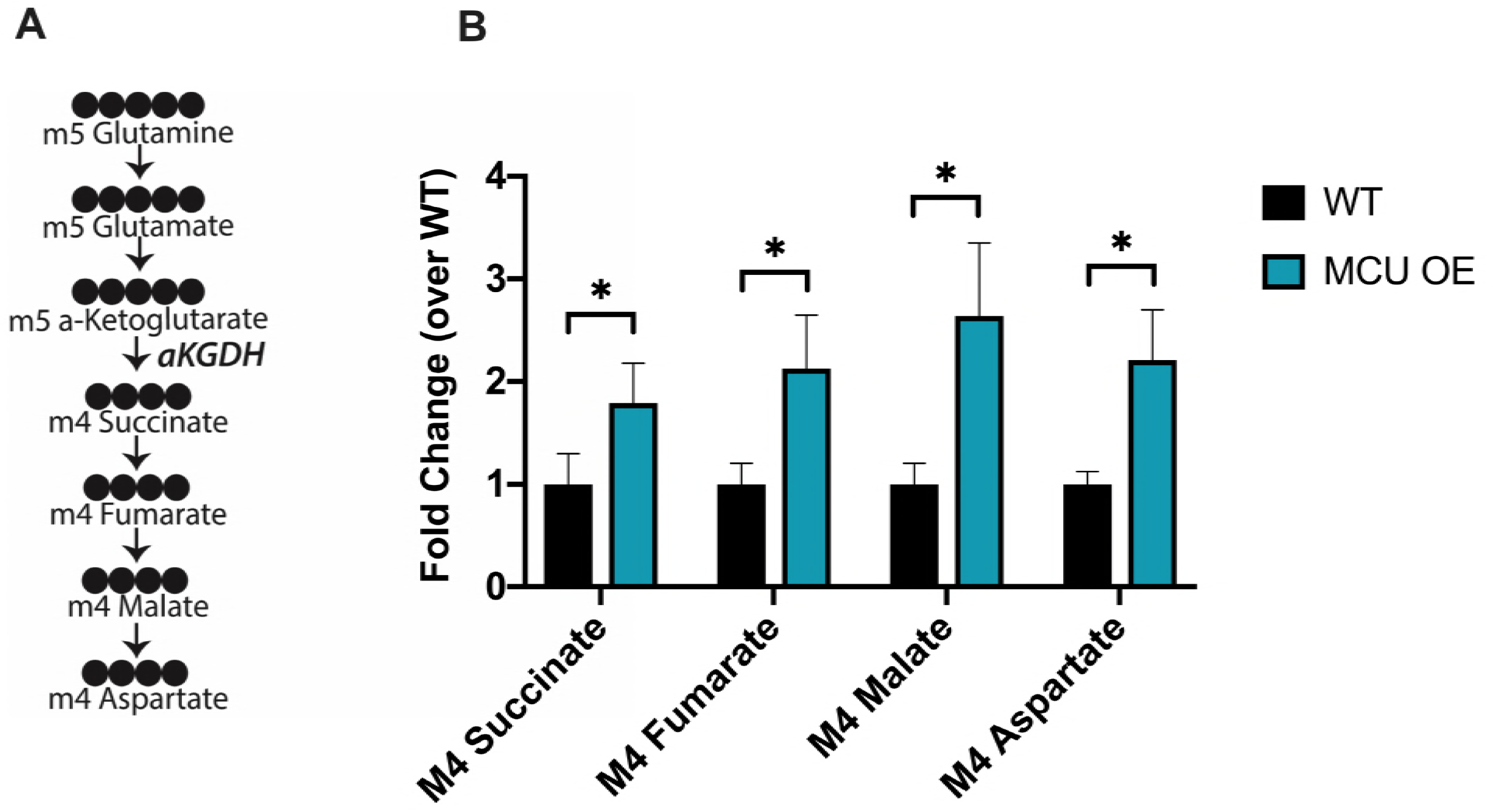
^13^C-Glutamine supplementation confirms increased α-KGDH activity in MCU OE retinas. A. Diagram showing how labelled carbons from U-^13^C-glutamine are incorporated into α-ketoglutarate and downstream metabolites. Shaded = labeled carbon, empty = unlabeled carbon. B. Levels of isotopomers in WT and MCU retinas supplied with 2 mM ^13^C-glutamine for 15 minutes. Data points represent averages from n=3 retinas from 3 different fish. * p < 0.05 using Welch’s t-test.

### MCU-overexpressing mitochondria are mislocalized and have abnormal morphology

In zebrafish cones, mitochondria are reported to be confined solely to the ellipsoid region (Tarboush et al., 2012). However, while imaging *gnat2*:mito-GCaMP3 larvae, we observed large mitochondrial clusters outside of this region in the MCU OE fish (**Figure 7A**). We recorded 8 hour time-lapses of live zebrafish larvae expressing *gnat2*:mito-GCaMP3 and *gnat2*:MCU-T2A-RFP. Cytosolic RFP was included to visualize cones. The timelapse recordings demonstrate directed movement of mitochondrial clusters away from the ellipsoid region toward the synapse (**Figure 7B, supplemental video 1**).

**Figure 7:**
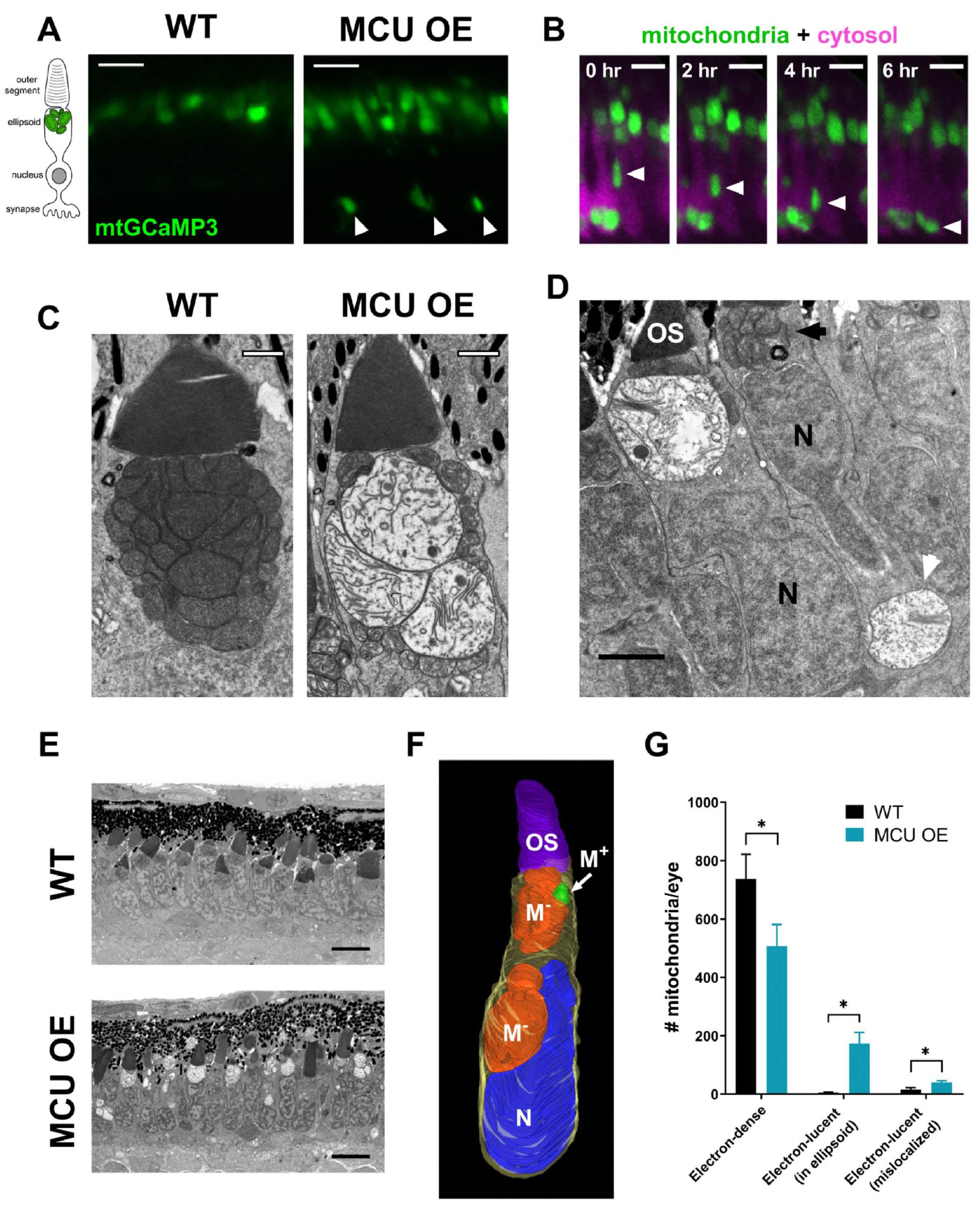
Mitochondria in MCU OE cones migrate away from the ellipsoid region and exhibit swelling, loss of electron-density, and disorganized cristae. A. Cone mitochondrial clusters in live larvae expressing *gnat2*:mtGCaMP3. WT mitochondria were imaged with higher laser settings to show localization. In MCU OE models, mitochondrial clusters were found near the synapse and nuclear layer (white arrows), which was not observed in WT siblings. Scale bar = 5 μm. B. Timelapse of a migrating mitochondrial cluster (green, white arrow) in live MCU OE *gnat2*:mtGCaMP3 larvae. Cone cell bodies express cytosolic RFP (magenta). MCU OE mitochondrial clusters slowly move from the ellipsoid region down to the synapse. Full video in supplemental data. Scale bar = 5 μm. C. EM images of cone mitochondria in WT sibling and MCU OE fish at 1 month of age. Mitochondria in MCU OE cones exhibit swelling, loss of electron density, and disorganized cristae. Scale bar = 1 μm. D. A single cone photoreceptor can contain both healthy mitochondria in the ellipsoid region (black arrow) and electron-lucent mitochondria near the synapse (white arrow). Electron micrograph from MCU larvae at 14 days of age. Scale bar = 2 μm. OS = outer segment, N = nucleus. E. Electron micrograph of MCU cones at 120 hours of age. Scale bar = 5 μm. F. 3D reconstruction of an MCU OE cone using serial block-face EM (synapse not shown). Electron-lucent mitochondria displace the nucleus of the cone to move toward the synapse region. OS = outer segment. M^−^ = electron-lucent mitochondria. M+ = healthy mitochondria. N = nucleus. Outline (yellow) = cell body. G. Quantification of cone mitochondrial phenotypes from EM images of whole zebrafish larval eyes (single slice at optic nerve) at 6 days of age. MCU OE cones have fewer “healthy” electron-dense mitochondria, and an increase in lucent mitochondria both in the ellipsoid and outside of this region. All mislocalized mitochondria observed had an electron-lucent phenotype. n=3 larvae for both WT and MCU OE fish. *p<0.05 using a t-test with the Holm-Sidak correction for multiple tests.

EM analysis reveals dramatic changes in mitochondrial structure in both mislocalized mitochondrial clusters and those in the ellipsoid region. Mitochondria in MCU OE cones increase in size to the scale of an entire normal mitochondrial cluster, they lose substantial electron density, and their cristae are disorganized (**Figure 7C**). This phenotype is heterogenous; some abnormal mitochondria have internal membrane stacks, clusters of electron dense material, or few internal membrane structures. Furthermore, many cones contain a mix of electron-dense and -lucent mitochondria; even cones containing mislocalized, electron-lucent mitochondria near the synapse can also contain what appear to be healthy mitochondria at the ellipsoid region (**Figure 7D**). This electron-lucent phenotype emerges soon after expression of MCU is first detectable at 78 hours post-fertilization and is widespread by 120 hours post-fertilization (**Figure 7E**). 3D reconstructions of MCU OE mitochondria suggest that movement of electron-lucent mitochondria away from the ellipsoid region is an active process, as these mitochondria deform the nucleus on their way toward the synapse (**Figure 7F, Supplemental Video 2**).

Using electron micrographs of whole larval eyes, we quantified mitochondria that were electron-lucent in the ellipsoid region and those that were mislocalized to the nuclear layer and synapse. With the caveat that the lucent mitochondria in MCU OE models are often much larger than their healthy counterparts, we found that relative to WT cones the MCU OE cones have fewer electron-dense mitochondria and substantially more electron-lucent mitochondria both within the ellipsoid region and outside of it (**Figure 7G**). Notably, electron-dense mitochondria were never observed outside of the ellipsoid region in either WT or MCU OE cones. WT cones rarely contained electron-lucent mitochondria (2.7 ± 0.6% of cone mitochondria); these were markedly smaller than lucent mitochondria in their MCU OE siblings (**Supplemental Figure 7A**), and most of these were mislocalized (2.0 ± 0.7% of cone mitochondria, **Supplemental Figure 7B**). Cones containing lucent and mislocalized mitochondria were more concentrated on the dorsal side of the retina, with fewer near the periphery of the retina where new photoreceptors form (**Supplemental Figure 7C**). Overall, we found that 37.6 ± 4.0% of cone ellipsoid clusters contained electron lucent mitochondria in MCU OE larvae (**Supplemental Figure 7D**). The widespread mitochondrial swelling and mislocalization phenotype of MCU OE fish is not specific to larval development and is retained as fish mature (**Supplemental Figure 7E**).

### Cones overexpressing MCU survive throughout early adulthood, but eventually degenerate

Despite the early emergence of mitochondrial abnormalities, we noted that larval cones otherwise remain intact and other retinal cell morphology is unperturbed (**Supplemental Figure 8A**). Fluorescent imaging of Tg(*gnat2*:GFP) fish showed that cones are conserved as zebrafish reach maturity at 3 months of age and throughout early adulthood despite their severely altered mitochondrial structure. However, some morphological disturbances (shorter cones and disorganized alignment) are apparent at these earlier ages. Severe cone degeneration eventually occurs later in adulthood; the cone number decreases, and many remaining cones round up and lose their outer segment integrity (**Figure 8A,B**). Rods remain intact and healthy rod outer segments persist even when cone loss is extensive (**Figure 8B, black arrowhead**).

**Figure 8:**
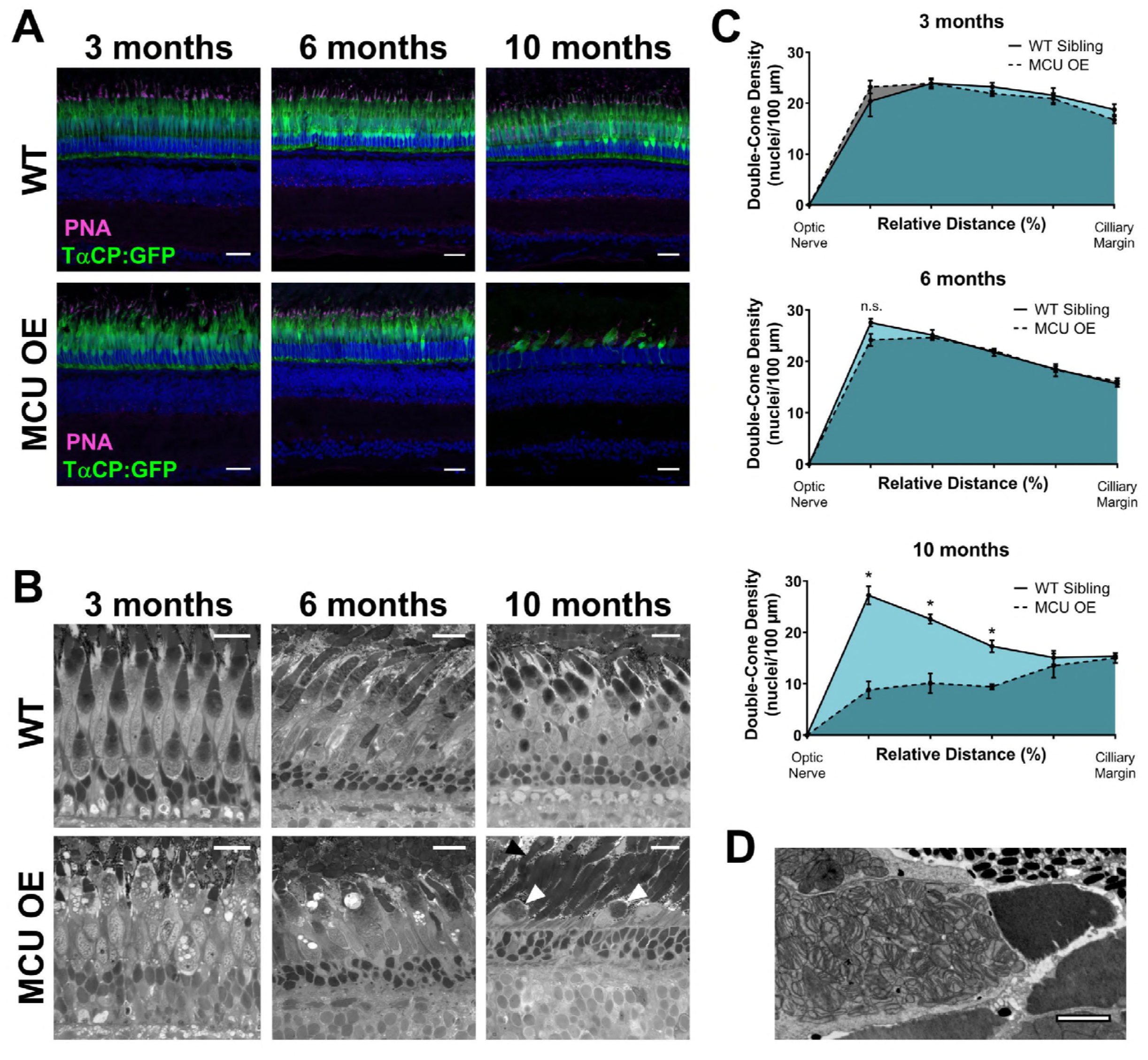
Cones overexpressing MCU survive well into early adulthood, but extensive loss is observed in late adulthood. A. WT sibling and MCU OE retinas stained with Hoescht (blue) and exhibiting fluorescence from *gnat2*:GFP in all cone types (green) at 3 months, 6 months, and 10 months of age. Cone outer segments are labelled with α-PNA (magenta). Scale bar = 25 μm. B. WT sibling and MCU OE retinas stained with Richardson’s stain at 3 months, 6 months, and 10 months of age. Swollen mitochondria can still be visualized in 3 month and 6 months cones. By 10 months, the cones are very few and have severe morphological disturbances (white arrow); however, rod mitochondria and outer segments remain intact (black arrow). Scale bar = 25 μm. C. Quantification of double-cone nuclei from the optic nerve to the ciliary margin in WT and MCU OE fish. Counts are an average of dorsal and ventral retinal slices in each fish, n = 3 WT and 3 MCU OE fish. *p<0.05 using a t-test with Holm-Sidak correction for multiple tests. D. Electron micrograph of fragmented mitochondria in a cone from 10-month-old MCU OE fish. We observed that many of the remaining cones at 10 months have severe mitochondrial fragmentation rather than the electron-lucent phenotype. Scale bar = 2 μm.

Quantification of double-cone nuclei, which are most easily distinguished from rod nuclei, revealed that cones in the MCU OE model are preserved at 3 months and 6 months, but severe cone loss occurs by 10 months (**Figure 8C, Supplemental Figure 8B**). Cone loss is consistently most apparent near the optic nerve; cones are only appreciably observed at the ciliary zone of the 10 month MCU OE zebrafish retina, where new cones form. Many of the remaining cones in the MCU OE model have severe mitochondrial fragmentation rather than large electron-lucent mitochondria (**Figure 8D**). Mitochondrial fragmentation is often associated with apoptosis (Frank et al., 2001; Lee et al., 2004; Youle and Karbowski, 2005). Cones in MCU OE models do not re-populate to WT density after 10 months, but rods remain intact and abundant (**Supplemental Figure 8C**). Notably, in 1 year old WT sibling retinas, there are multiple cones with large, electron-lucent mitochondria in the clusters; these were similar in appearance to MCU OE electron-lucent mitochondria (**Supplemental Figure 8D**).

## Discussion

Photoreceptors in normal human retinas are remarkable in their ability to survive the entire human life span. Although perturbations to Ca^2+^ homeostasis are implicated in many degenerative diseases, normal photoreceptors can withstand the large intracellular Ca^2+^ fluctuations that occur during normal photoreceptor function. Identification of the MCU protein has allowed us to investigate its role in a biological context; the abundance of direct and indirect regulators of MCU and variability in expression of these regulators across tissues necessitates empirical, cell-type-specific study. Here we investigated the influence of MCU expression on cone photoreceptor function and survival. Our findings demonstrate an important role of mitochondrial Ca^2+^ uptake in regulating functional and structural aspects of cone photoreceptors. This work also reveals a surprising resistance of photoreceptors to the stress caused by elevated mitochondrial matrix Ca^2+^. Our results indicate that in photoreceptors, homeostatic adaptive responses to high matrix Ca^2+^ are robust.

### Effect of MCU OE on Ca^2+^ homeostasis and the functional consequences

Our findings suggest that MCU expression is a limiting factor in basal cone mitochondrial Ca^2+^ uptake, as overexpression of MCU increases the resting concentration of free Ca^2+^ in the mitochondrial matrix. EMRE, required for Ca^2+^ uptake via MCU in vertebrates, does not seem to be limiting in cones; this is consistent with a recent study demonstrating that EMRE is produced in excess and degraded when MCU is not present (Sancak et al., 2013, Tsai et al., 2017). Enhancing mitochondrial Ca^2+^ uptake in cones reduces the amplitude and duration of Ca^2+^ transients in the cell body, where mitochondria reside. However, only partial restoration to WT levels can be achieved in the MCU OE model by pharmacologically inhibiting MCU. This could be due to factors other than mitochondrial Ca^2+^ uptake being affected in the MCU OE model (such as ER Ca^2+^ levels), insufficient permeability of Ru360 into cells, or an abundance of MICU1/MICU3 which can block Ru360 binding to MCU (Paillard et al., 2018).

Extrusion of Ca^2+^ from the outer segment of photoreceptors is thought to be accomplished primarily by plasma membrane Na+/Ca^2+^, K+ exchangers NCKX1 in rods, and NCKX2 and NCKX4 in cones (for recent review, see Vinberg et al., 2018). However, *Nckx1^−/−^* rods appear to clear cytosolic Ca^2+^ through a slower, yet unidentified mechanism (Vinberg et al., 2015a). Moreover, cones that are deficient of both NCKX2 and NCKX4 can respond to light, light-adapt, and degenerate rather slowly, suggesting that there is an additional NCKX-independent pathway that clears Ca^2+^ from cone outer segments (Vinberg et al., 2017). The faster clearance of cytosolic Ca^2+^ by mitochondria in MCU OE cones is accompanied by accelerated photoresponse recovery, consistent with faster Ca^2+^ clearance. These results suggest that MCU may be the NCKX-independent mechanism that contributes to the clearance of Ca^2+^ from the cone outer segment.

Increased light sensitivity of MCU OE cones (**Supplemental Figure 5C**) is surprising considering that the inactivation of cone phototransduction was faster in these cones. The gain of cone phototransduction activation reactions, normally thought to be Ca^2+^-independent, is somehow increased by altered Ca^2+^ homeostasis due to MCU overexpression. We also found that the maximal light response amplitude was decreased in MCU OE cones (**Supplemental Figure 5B**), possibly due to misalignment or loss of cones at the age tested (7 months). Altogether, these functional changes demonstrate that mitochondrial Ca^2+^ uptake in photoreceptors can modulate the kinetics of the photoreceptor response to light, independent of neurotransmitter release and downstream responses. It is likely that this phenomenon is not restricted to zebrafish or cones, as human patients with mtDNA diseases have delayed recovery of the rod photoreceptor response (Cooper et al., 2002).

### Effect of MCU OE on mitochondrial metabolism in the retina

Since photoreceptors experience the highest levels of intracellular Ca^2+^ in darkness when O2 consumption is highest, Ca^2+^ could play an important role in stimulating increased TCA cycle activity (Okawa Fain 2008, Krizaj Copenhagen 2002). We have previously observed that photoreceptors have higher α-KGDH activity in darkness (Du Rountree 2015). Here, we observe that both IDH and α-KGDH are stimulated by increasing Ca^2+^ in cone mitochondria, which supports this hypothesis.

An increase in the P-PDH/total PDH ratio is a common metabolic phenotype in MCU KO tissues, so we initially hypothesized that increasing mitochondrial Ca^2+^ would decrease the P-PDH/total PDH ratio by stimulating PDP1c (for a review of metabolic phenotypes in MCU KO tissues see Mammucari, Raffaello 2018). However, we found that this was not the case. Since the P-PDH/total PDH ratio also does not decrease when MCU is overexpressed in muscle cells, it is possible that changes in mitochondrial bioenergetics resulting from increased mitochondrial Ca^2+^ feed into the complex regulation of PDH (Mammucari, Gherardi 2015). For example, stimulation of α-KGDH and IDH activity may result in higher NADH levels in MCU OE cones, which in turn stimulates PDH kinase to balance increased PDP1c activity.

### Effect of MCU OE on mitochondrial homeostasis and subsequent cell death

While some of our results align with predicted consequences of increased mitochondrial Ca^2+^ uptake, our work also revealed some surprising observations. The most unexpected finding in MCU OE models was the presence and selective movement of electron-lucent mitochondria away from the ellipsoid region of cones toward the synapse. We also occasionally observed electron-lucent structures that appeared to be mislocalized mitochondria in WT fish, especially in older animals (**Supplemental 7A,B; Supplemental 8D**). We hypothesize that this mechanism of mitochondrial sorting and movement occurs in normal cones, but it is more active in MCU OE models due to extensive mitochondrial disruption. This hypothesis is strengthened by recent work with the mito-QC mouse showing photoreceptor mitochondria that were being recycled by lysosomes were specifically present in the outer nuclear layer, away from the ellipsoid region (McWilliams et al., 2016). Additionally, mitochondria in rod-specific Nrf1 KO mice were observed near the outer limiting membrane of rods, away from the ellipsoid region (Kiyama et al., 2018). Many of the models of mitochondrial localization focus on how cytosolic Ca^2+^ changes predicate mitochondrial movement, concentrating mitochondria in high Ca^2+^ regions (Barnhart, 2016; Chen and Sheng, 2013; MacAskill et al., 2009; Wang and Schwarz, 2009). However, mitochondrial matrix Ca^2+^ and interactions between MCU and the transport protein Miro1 may also play a role in mitochondrial movement (Chang et al., 2011; Niescier et al., 2013, 2018). These studies, along with our findings, highlight the need for further investigation of the role of mitochondrial Ca^2+^ content and subsequent stress in selective, directed mitochondrial movement.

We initially suspected that the swollen, electron-lucent mitochondria in MCU OE cones were a manifestation of Ca^2+^ overload that would subsequently trigger apoptosis and rapid loss of cones. Mitochondria undergoing the permeability transition that precedes apoptosis exhibit excessive osmotic swelling and loss of electron density similar to the changes that are observed in MCU OE cones (Elustondo et al., 2016; Hunter et al., 1959; Zoratti and Szabò, 1995). Remarkably, MCU OE cones survive until late adulthood despite their abnormal morphology and electron-lucent mitochondria. This finding suggests that cones are particularly resistant to mitochondrial stress that would trigger apoptosis in other cell types. Furthermore, the occasional presence of large, electron-lucent mitochondria in WT retinas as the zebrafish mature (**Supplemental Figure 8D**) suggests that this phenotype may even be a symptom of cumulative mitochondrial stress in cones as they age. This phenomenon is unlikely to be specific to zebrafish cones, as large, electron-lucent mitochondria also have been observed in aging cones in human retinas (Nag and Wadhwa, 2016). Increased resistance to mitochondrial stress may be an adaptation necessary for cones, which need to meet excessively high energetic demands throughout a lifetime.

### Future Directions, Limitations, and Broader Impact

Despite the network of regulatory proteins that influence Ca^2+^ flux through MCU, solely altering MCU protein levels can influence mitochondrial Ca^2+^ and cell function. Overexpression of MCU in cortical neurons exacerbates excitotoxic cell death (Qiu 2013). However, overexpression of MCU does not always result in negative consequences. Overexpressing MCU in muscle cells increases their size and protects them from denervation-induced atrophy, and increasing MCU expression in diabetic heart models improves cardiac function (Mammucari et al., 2015; Suarez et al., 2018). MCU activity varies significantly across tissues, as does protein expression of MCU and its regulators (Fieni et al., 2012; Plovanich et al., 2013; Raffaello et al., 2013). It is likely that the consequences of MCU overexpression depend on the presence of its regulators. The MICU1:MCU ratio appears to be critical in regulating the mitochondrial response to Ca^2+^ transients; muscles, with the lowest MICU1:MCU ratio of characterized tissues, notably benefit from MCU overexpression (Mammucari et al., 2015; Paillard et al., 2017). Our qPCR results suggest that the retina has high ratios of MICU1:MCU and MICU3:MCU compared to other neural tissues, which would imply that the limited MCU channels present in the retina are primed for highly efficient Ca^2+^ uptake via a MICU1-MICU3 dimer. This could explain why MCU overexpression causes such massive perturbations in cones compared to other tissues. Future studies will characterize the functional roles of MICU1 and MICU3 in retinal cells.

A key finding from our study is that cones are highly tolerant of mitochondrial stress. 100-fold overexpression of MCU is far above the amount of MCU normally present in retina. It is possible that some of the phenotypes of the MCU OE model are secondary effects of excess protein, such as stress on the ubiquitin-proteasomal system. However, the effects of MCU OE on cytosolic Ca^2+^ and the citric acid cycle are consistent with previous studies on mitochondrial Ca^2+^, suggesting many of the phenotypes observed were linked to increased mitochondrial Ca^2+^ uptake.

Overall, our work supports a model in which cone photoreceptors maintain low MCU to fine-tune the many changes associated with increased mitochondrial matrix Ca^2+^ and protect from Ca^2+^-associated mitochondrial disruption. The cone MCU OE model can also be used in future studies to elucidate a mechanistic explanation behind selective sorting of abnormal mitochondria and the adaptations triggered to maintain viability and functionality despite chronic stress. Even more broadly, the distinct localization of mitochondria in zebrafish cones makes the MCU OE model an attractive system to study basic cellular processes underlying mitochondrial movement.

## Materials and Methods

### Zebrafish Maintenance

Experiments with zebrafish were authorized by the University of Washington and University of Utah Institutional Animal Care and Use Committees. All fish used in this analysis were maintained in the University of Washington South Lake Union aquatics facility or the Centralized Zebrafish Animal Resource (CZAR) at the University of Utah at 27.5C on a 14 h/10 h light/dark cycle, and were maintained in the Roy^−/−^ genetic background. All wild-type fish (WT) used in analysis were age-matched siblings to Tg(gnat2:MCU-T2A-RFP) fish (MCU OE) or age-matched siblings to pde6c^w59^ (pde6c^−/−^) (Stearns et al., 2007). Fish used for slice preparation, protein quantification, and metabolomics analysis were male and female siblings between 3 and 6 months of age. Fish used in ERG analysis were male and female siblings collected at 7 months of age. For histological analysis, ages of sibling fish are included in the figure and legend.

### Zebrafish MCU antibody

The cDNA encoding amino acids 21 – 202 of *Danio rerio* MCU (NM_001077325) was cloned downstream of GST using the pGEX-2T (GE) expression vector. Overexpression was induced in *E.coli* (BL21) by addition of 1mM IPTG at 0.2 OD followed by incubation with vigorous shaking for 5h at 37° C. The tiny fraction of soluble fusion was purified using glutathione sepharose following the manufacturer’s instructions (GE Healthcare). Polyclonal antibodies were generated using injections of 0.5 – 1mg protein (R and R Research Co.). Two columns were used to clean the serum. One column contained total *E.coli* proteins covalently coupled to cyanogen bromide beads and the second column contained purified GST protein coupled to cyanogen bromide beads (GE Healthcare). Serum was cleaned by sequential incubations of 3 – 5 hours at room temperature with each column after which it was analyzed on an SDS page gel for lack of cross reactivity with GST and *E.coli* proteins from a total cell extract. Identification of MCU was validated by the absence of a protein of the correct molecular weight in extracts obtained from a CRISPR generated KO strain. The zebrafish MCU antibody was used at a dilution of 1:750 for western blotting and 1:50 for immunohistochemistry.

### Zebrafish models

The transgenic zebrafish lines Tg(gnat2:GCaMP3), Tg(gnat2:EGFP), and Tg(mito-GCaMP3) have been described previously (Giarmarco et al., 2017; Kennedy et al., 2007; Ma et al., 2013). Generation of the global MCU KO line was performed using gRNA with the following sequence 5’-CCTCATACCTGGTGCAGCCCCCC-3’ using methods as previously described (Brockerhoff, 2017). For generation of the Tg(*gnat2*:MCU-T2A-RFP) line, zebrafish MCU cDNA was isolated from WT zebrafish larvae (5 dpf) using the forward primer 5’-AGAGATGGCTGCGAAAAGTGT-3’ and reverse primer 5’-TTCTCATCAGTCCTTGCTGGT-3’. Overhang qPCR methods in conjunction with Fast-Cloning were used to add the T2A ribosomal stalling sequence and the RFP protein coding sequence; this was cloned into a pCR8/GW vector (Invitrogen) (Li et al., 2011). Plasmids were assembled using the Gateway-Tol2 system (Villefranc et al., 2007). Expression of MCU-T2A-RFP was driven by the cone transducin alpha promoter (TαCP, gnat2), and the RFP coding sequence was flanked by a polyA tail sequence to increase transcript stability (Kennedy et al., 2007). A destination vector with a sBFP2 heart marker for aid in transgenic identification was obtained from Cecilia Moens (Kremers et al., 2007). The fully assembled construct was injected into embryos at the 1-cell stage with Tol2 transposase mRNA. Larvae mosaic for the transgene were raised to adulthood to identify founder carriers. A single F_0_ founder was used to generate F_1_ fish that were screened for a single insertion of the transgene; F_2_ fish from two F_1_ substrains with a single insertion were used for analysis in this study.

### Primers for qRT-PCR

All designed primers were empirically tested to confirm primer efficiency was between 90-110%. Only primers passing this benchmark were used for analysis. Primer sequences for the reference genes *EF1a, b2m, Rpl13a*, and *TBF* were identical to previous reports testing zebrafish reference gene stability *(EF1a, Rpl13a*: (Tang et al., 2007), *b2m, TBP*: (McCurley et al., 2008)).

Elongation Factor 1 alpha (EF1a):

Forward: 5’-CTGGAGGCCAGCTCAAACAT-3’

Reverse: 5’-ATCAAGAAGAGTAGTACCGCTAGCATTAC-3’

Beta-2-microglobulin (b2m):

Forward: 5’-GCCTTCACCCCAGAGAAAGG-3’

Reverse: 5’-GCGGTTGGGATTTACATGTTG-3’

Ribosomal protein L13a (Rpl13a):

Forward: 5’-TCTGGAGGACTGTAAGAGGTATGC-3’

Reverse: 5’-AGACGCACAATCTTGAGAGCAG-3’

TATA-box-binding protein (TBP):

Forward: 5’-CGGTGGATCCTGCGAATTA-3’

Reverse: 5’-TGACAGGTTATGAAGCAAAACAACA-3’

MICU1:

Forward: 5’-ACGTTAAAGCAGAATCGTAGAGG-3’

Reverse: 5’-CGCAAGCGGTACATATCAGAC-3’

MICU2:

Forward: 5’-ACTGAGTACCTGTTTCTCCTCAC-3’

Reverse: 5’-GGTCCATTTACTTTCTTCAGCTTCT-3’

MICU3a:

Forward: 5’-CGTCCCATGAGCATCGTTTC-3’

Reverse: 5’-TCCAACTCCTGTTTGGTGAGG-3’

MICU3b:

Forward: 5’-GCTTGGTGCAAGAATAGTTCTCTTT-3’

Reverse: 5’-TGCAGGTTGTCCATGAATCTGT-3’

### qRT-PCR

An Applied Biosystems 7500 Fast Real-Time PCR System in conjunction with iTaq™ Universal SYBR® Green Supermix (Bio-Rad, 1725120) was used for qPCR measurements according to the manufacturer’s instructions. The reference genes *EF1a, b2m, TBF*, and *rpl13a* were screened across the tissue panel using NormFinder to identify reference genes with the highest stability (Andersen et al., 2004). NormFinder identified the combination of *EF1a* and *b2m* as most stable for retina-brain comparisons and *EF1a* as most stable for retina-heart comparisons. Quantification of relative mRNA quantity used 3 biological replicates of each tissue, each performed in technical triplicate. From each technical triplicate, the average C_t_ value for the gene of interest and reference gene(s) were used to generate a ΔC_t_ value for each biological replicate. Comparing each tissue of interest to the retina generated a ΔΔC_t_ value; these were converted to a normalized expression level using the 2^−ΔΔCt^ method (Livak assumptions). Standard error of the ΔC_t_ value for each tissue was propagated to the final comparison using standard error propagation rules. Calculations were based off the geNorm method of qPCR normalization (Vandesompele et al., 2002).

### Commercial antibodies and stains

Mitochondrial cytochrome oxidase, MTCO1 (Abcam, ab14705, RRID:AB_2084810). Used in IHC and immunoblotting at 1:1000 dilution.

Succinate dehydrogenase B, SDHB (Abcam, ab14714, RRID:AB_301432).

Used in IHC and immunoblotting at 1:1000 dilution.

Pyruvate Dehydrogenase E1 subunit, PDH (Abcam, ab110334, RRID:AB_10866116) Used in immunoblotting at 1:1000 dilution.

Phosphorylated Pyruvate Dehydrogenase E1 subunit Ser293, P-PDH (EMD Millipore, ABS204, RRID:AB_11205754)

Used in immunoblotting at 1:2000 dilution.

Pyruvate Kinase, PK (Abcam, ab137791)

Used in immunoblotting at 1:1000 dilution.

Hoechst 33342, Trihydrochloride, Trihydrate stain (ThermoFischer, H3570).

Used in IHC at 5 μM concentration.

Lectin PNA Alexa Fluor 647 conjugate (ThermoFischer, L32460).

Used in IHC at 1:200 dilution after suspending at a concentration of 1 mg/mL in H2O.

Goat Anti-Mouse IgG H&L, Alexa Fluor 488 (Abcam, ab150113, RRID:AB_2576208).

Used in IHC at 1:1000 dilution.

Goat Anti-Rabbit IgG H&L, Alexa Fluor 647 (Abcam, ab150083, RRID:AB_2714032).

Used in IHC at 1:1000 dilution.

IRDye 800CW donkey anti-rabbit IgG (H+L) (LI-COR Biosciences, 925-32213, RRID: AB_2715510).

Used at 1:5000 dilution for immunoblotting.

IRDye 680RD donkey anti-mouse IgG (H+L) (LI-COR Biosciences, 925-32212, RRID: AB_2716622).

Used at 1:5000 dilution for immunoblotting.

IRDye 680RD donkey anti-rabbit IgG (H+L) (LI-COR Biosciences, 925-68073, RRID:AB_2716687).

Used at 1:5000 dilution for immunoblotting.

IRDye 800CW goat anti-mouse IgG (H+L) (LI-COR Biosciences, 925-32210, RRID:AB_2687825).

Used at 1:5000 dilution for immunoblotting.

### Mitochondrial enrichment and sample preparation for immunoblotting

Organs were snap-frozen in liquid nitrogen after collection, then homogenized with a dounce homogenizer in 50 mM Tris buffer containing sucrose (200 mM), NaCl (150 mM), and EGTA (1 mM) with a protease inhibitor mini tablet (ThermoFischer, 88666). Homogenized samples were centrifuged at a low speed of 1,000 x g for 10 minutes at 4°C, then the supernatant (containing mitochondria) was collected and centrifuged at a high speed of 17,000 x g for 45 minutes. The supernatant was discarded and the pellet (containing mitochondria) was homogenized for 1 minute in RIPA buffer. Homogenized mitochondria were sonicated on ice for three 5 second pulses. A standard BSA assay using Pierce™ BCA Protein Assay Kit (ThermoFischer, 23225) was performed according to the manufacturer’s instructions for protein concentration determination. Samples were diluted with RIPA buffer to ensure an equal volume and equal protein concentration of each sample could be loaded into wells for immunoblotting.

### Immunoblotting

Samples were loaded into wells on 12-14% acrylamide gels made in-house. Each sample contained 20% 5X SDS buffer containing β-mercaptoethanol. After running the gel at 150V for 1 hr, gels were transferred onto PVDF membranes (Millipore, IPFL00010) and blocked for 1 hr at room temperature in LI-COR Odyssey Blocking Buffer (LI-COR, 927-40000). Primary antibodies were diluted in blocking buffer at specified concentrations and incubated overnight at 4°C. Membranes were washed with PBST and PBS, then incubated with secondary antibody for 1 hr at 25°C and washed again before imaging. Membranes were imaged and bands were quantified using the LI-COR Odyssey CLx Imaging System (RRID:SCR_014579).

### Immunohistochemistry (IHC) and degeneration quantification

All adult eyes were isolated from light-adapted zebrafish, and a small incision in the cornea was made to allow 4% paraformaldehyde fixative to enter the eye. Whole larvae were euthanized then incubated in 4% paraformaldehyde. After fixation overnight at 4°C, eyes were rinsed in PBS then subject to a sucrose gradient (20% and 30%), embedding in OCT, and cryosectioned at 12 μm. For sections stained with MCU antibody, antigen retrieval was performed by steaming sections in 10 mM sodium citrate (0.05% Tween-20, pH 6.0). Sections were washed in PBS, then blocked in PBS containing 5% donkey serum, 2 mg/mL bovine serum albumin, and 0.3% Triton X-100 for 1 hr. Primary antibodies were diluted in this buffer as specified, then applied to cryosection overnight at 4°C. Secondary antibodies were diluted as specified and applied to section for 1 hr in darkness at 25C. For PNA-labelled samples, sections were incubated in diluted PNA-647 for 30 minutes at 25°C. Tissues were washed, incubated in Hoescht stain for 10 minutes, then mounted in Fluoromount-G® (SouthernBiotech, 0100-01) under glass coverslips. Slides were imaged using a Leica LSP8 confocal microscope with a 63X oil objective. Leica LAS-X software (RRID:SCR_013673) was used to acquire images.

For quantification of cone nuclei in *gnat2*:GFP fish, high-resolution images of whole zebrafish retina slices were stitched together using ImageJ Grid/Collection stitching (Preibisch et al., 2009). Both the dorsal and ventral regions of the retina were straightened along the cone nuclei axis using ImageJ from the optic nerve to the cilliary margin. This axis was divided into 5 equal parts, then double-cone nuclei were counted in each region, normalizing to the length in μm (height of the region was equal across samples, double cone nuclei are along a single axis). GFP was used to confirm that the double-cone nuclei counted were indeed cone nuclei. All counting was performed blinded (masked) to sample identity.

### Live larval imaging of mtGCaMP3

Larvae used for imaging were maintained in embryo media containing 0.003% 1-phenyl 2-thiourea (PTU, Sigma-Aldrich P7629) starting at 20 hours post-fertilization. Live zebrafish larvae were analyzed at 6 days postfertilization (dpf) by transferring to 0.5% low melting point agarose containing embryo media with 0.003% PTU and 0.02% (w/v) Tricaine (Sigma-Aldrich, E10521). Larvae were positioned in agarose in a petri dish containing embryo media and 0.02% (w/v) tricaine to prevent drying out. Imaging of slices was performed using an Olympus FV1000 in conjunction with Olympus FluoView FV10-ASW software (RRID:SCR_014215). A 40X water objective was used for imaging. The excitation/emission wavelengths used for mito-GCaMP3 were 488/510 nm. Timelapse images of live larvae were collected with a z-depth of 2 μm and were collected every 20 minutes. Images of total eye mitochondrial clusters were also collected at a z-depth of 2 μm. For quantification of total mito-GCaMP3 fluorescence, images of whole larval eyes were collected, and a fixed ROI centered on the nasal region of the retina was used for quantification. This region near the ventronasal patch is comprised of the most mature cone photoreceptors (Schmitt and Dowling, 1999).

### Retinal slice imaging of GCaMP3 and mito-GCaMP3

Slices were prepared as described previously (Giarmarco et al., 2017, 2018). For GCaMP3 cytosolic clearance experiments, slides were preincubated in KRB containing 0 mM Ca^2+^ and 0.4 mM EGTA for 10 minutes. Images of single optical slices were collected every 2 seconds. A 5 mM CaCl_2_ bolus (accounting for EGTA) was injected into the slice imaging chamber 30s after the initial timelapse collection to establish baseline fluorescence. These experiments in the MCU-overexpressing fish were additionally performed in the presence of Ru360 (Millipore, 557440) at 100 μM, in which slices were incubated for 1 hr prior to incubation in 0 mM Ca^2+^ media. Retinas treated with Ru360 were maintained in Ru360 throughout timelapse experiments. Dying cells near the cut edge that were constitutively loaded with Ca^2+^ and cells that did not respond to Ca^2+^ were not included in analysis.

For mito-GCaMP3 timelapse experiments, z-stacks of 15, 2 μm slices were collected every 30 seconds. Retinal slices in modified KRB containing 2 mM CaCl_2_ were first imaged for 5 minutes to establish baseline mito-GCaMP3 fluorescence. Next, the chamber was injected with ionomycin (Sigma, 407950) to a final concentration of 5 μM (prepared in DMSO, at a final concentration of 0.1%) for another 5 minutes of image collection. Finally, an excess of EGTA (5 mM) was injected to chelate the 2 mM Ca^2+^ present in solution and images were collected for another 5 minutes. Dying cells containing fragmented mitochondrial clusters constitutively loaded with Ca^2+^ and clusters that did not respond to ionomycin were not included in analysis. Additionally, any clusters where the maximum fluorescence signal in the presence of ionomycin was completely saturated were excluded from analysis.

The excitation/emission wavelengths used for both GCaMP3 and mito-GCaMP3 were 488/510 nm. Timelapses were analyzed using ImageJ + Fiji software (SCR_002285). Images were corrected for X-Y drift using the MultiStackReg plugin of ImageJ. For both cell body GCaMP3 and mito-GCaMP3 fluorescence ex vivo timelapses, fixed ROIs were used to quantify average fluorescence signal across the cluster/cell at every time point. Fluorescence of cytosolic GCaMP3 for timelapse analysis are reported as F/F_0_, where F_0_ is the baseline fluorescence. For mito-GCaMP3 fluorescence, the relative fluorescence at maximum was set to 100% for normalization. We used the equation 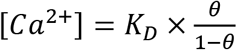, where 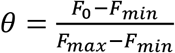 to approximate [Ca^2+^]_mito_, where F_0_ is the average “baseline” fluorescence, F_max_ is maximum fluorescence upon ionomycin addition, and F_min_ is the baseline fluorescence upon EGTA addition. We approximated the K_D_ of GCaMP3 at 345 nM (from Chen et al., 2013) for the calculation.

### Electroretinograms (ERG)

Zebrafish were briefly dark-adapted (~30min), before euthanization by icewater immersion. Eyes were enucleated into Modified Salamander Ringer’s solution(mM): NaCl 110, KCl 2.5, CaCl_2_ 1.0, MgCl_2_ 1.6, HEPES 10.0, Glucose 10.0, with pH adjusted to 7.8 with NaOH. The eyes were hemisected and retina isolated from the eyecup. All procedures after the dark adaptation were performed under dim red light. *Ex vivo* Electroretinogram (ERG) recordings were performed as described previously (Vinberg and Kefalov, 2015b). Isolated retinas were mounted photoreceptor side up onto the specimen holder (Vinberg et al., 2014), and perfused with Modified Salamander Ringer’s solution, supplemented with 40μM DL-AP4 (Tocris Bioscience) and 40μM CNQX (Tocris Bioscience) to isolate the photoreceptor component of the ERG signal (A-wave). The rate of perfusion was ~5 ml/min. and the experiments were conducted at room temperature (~23 °C).

ERG signal was first amplified (100X) and low-pass filtered at 300 Hz by a differential amplifier (DP-311, Warner Instruments), and data was acquired at 10KHz using a Sutter IPA amplifier/digitizer (Sutter Instrument, CA). A High-Power LED light source (Solis-3C, Thorlabs, Newton, NJ), with filter for red light (630 nm, Semrock, Rochester, NY) and LED driver (DC2200, Thorlabs) were used to provide the flashes of light stimuli, durations ranged from 5-100ms. The SutterPatch software (SutterPatch v1.1.2, Sutter Instrument, CA) drove both stimulus generation and data acquisition via the IPA amplifier’s analogue output and input, respectively. Light stimuli were calibrated before experiments using a calibrated photodiode (FDS100-CAL, Thorlabs, Newton, NJ) and flash intensities converted to photon/μm^2^.

Data analysis, including statistical analysis and figure preparation, was performed with GraphPad v 8.00 (for Windows, GraphPad Software, CA, USA). Normalized responses were calculated for each retina by dividing the response amplitude data by the maximal amplitude measured at the peak/plateau of the response to the brightest flash.

### Metabolic flux analysis

Krebs-Ringer bicarbonate (KRB) buffer optimized for flux analysis (Du, Linton, Hurley 2013a) was used in these experiments. Zebrafish retinas were first dissected in KRB buffer containing U-^12^C-glucose or U-^12^C-glutamine at the same concentration they would be incubated in. After dissection, retinas were placed in dishes of pre-warmed KRB containing either U-^13^C glucose (5 mM, Cambridge Isotopes, CLM-1396) or U-^13^C glutamine (0.1 - 2 mM, Cambridge Isotopes, CLM-1822). Retinas were incubated in this solution for the specified time points at 28°C in a NAPCO Series 8000 WT CO_2_ incubator (5% CO_2_), then washed in ice-cold PBS and flash frozen in liquid nitrogen. Metabolites from each time point were extracted using ice-cold 80% MeOH and lyophilized. Two-step derivatization was performed by the addition of 20 mg/mL Methoxyamine HCl dissolved in pyridine, followed by *tert*-butyldimethylsilyl. Metabolites were analyzed on an Agilent 7890/5975C GC-MS as described extensively in previous work (Du et al., 2013a, 2013b, 2015, 2016). Metabolic flux experiments were repeated a minimum of twice, using three retinas from three different zebrafish for each condition in each experiment. Data shown are results from one representative experiment.

### Electron microscopy and Richardson’s staining

Adult zebrafish eyes were enucleated and a small incision was made in the cornea to allow fixative (4% glutaraldehyde in 0.1 M sodium cacodylate buffer, pH 7.2) to enter the eye. Tissues were stored at 4°C before postfixation in osmium ferrocyanide (2% osmium tetroxide/3% potassium ferrocyanide in buffer) for 1 hr, followed by incubation in 1% thiocarbohydrazide for 20 min. Samples were then incubated in 2% osmium tetroxide for 30 min at RT, and stained with 1% aqueous uranyl acetate overnight at 4°C. Samples were next stained en bloc with Walton’s lead aspartate for 30 min at 60°C, dehydrated in a graded ethanol series, and embedded in Durcupan resin. Sections of tissue were cut at 60 nm thickness and imaged using a JEOL JEM-1230 transmission electron microscope or Zeiss Sigma VP scanning electron microscope. Samples of larval zebrafish eyes were imaged in conjunction with a Gatan 3View2XP ultamicrotome apparatus to generate stacks of EM images, which were aligned using TrakEM2 software (RRID:SCR_008954). Position in the eye for EM imaging was confirmed by cutting slices of tissue and staining with Richardson’s stain (Richardson et al., 1960). These slices were imaged for histological analysis using a Nikon Eclipse E1000 with a Nikon Plan Apo 100X/1.40 DIC lens. Nikon ACT-1 software was used for image capture.

### Statistics

Numerical results in text are reported as mean ± standard error of the mean unless otherwise stated. Statistical tests were performed using Graphpad Prism v 8.00 software. For statistical analysis, replicates (n) were always defined as biological replicates. Information on what constitutes n (e.g. larvae, retinas, cells) is listed in the figure legend of each experiment. Samples sizes were estimated based on previous experiments (Sakurai et al., 2015; Giarmarco et al., 2017; Du et al., 2016). For data sets with sufficient n to analyze population distribution, tests for normality were administered (Anderson-Darling, D’Agostino & Pearson, Shapiro-Wilk, Kolmogorov-Smirnov). For data sets that did not pass a majority of normality tests, the median is instead reported along with the interquartile range (Q1 and Q3).

## Acknowledgements

We thank Stanley Kim, Jeanot Muster, and Ashlee Evans for assistance in zebrafish husbandry and maintenance at the University of Washington South Lake Union aquatics facility (ISCRM Aquatics Facility). We would also like to acknowledge the Centralized Zebrafish Animal Resource (CZAR) at the University of Utah for providing zebrafish husbandry, laboratory space, and equipment to carry out portions of this research. Expansion of the CZAR is supported in part by NIH grant # 1G20OD018369-01. This work was supported by PHS NRSA T32GM007270 from NIGMS (RAH); NIH grants NEI EY026020 (JBH and SEB), NEI EY028645 (SEB) and P30EY001730 (UW Vision Core). This work was additionally supported by NIH EY014800 (John A. Moran Eye Center), and an Unrestricted Grant from Research to Prevent Blindness, New York, NY, to the Department of Ophthalmology & Visual Sciences, University of Utah.

## Competing Interests

The authors declare no competing financial interests.

---

**Supplemental Figure 4:**
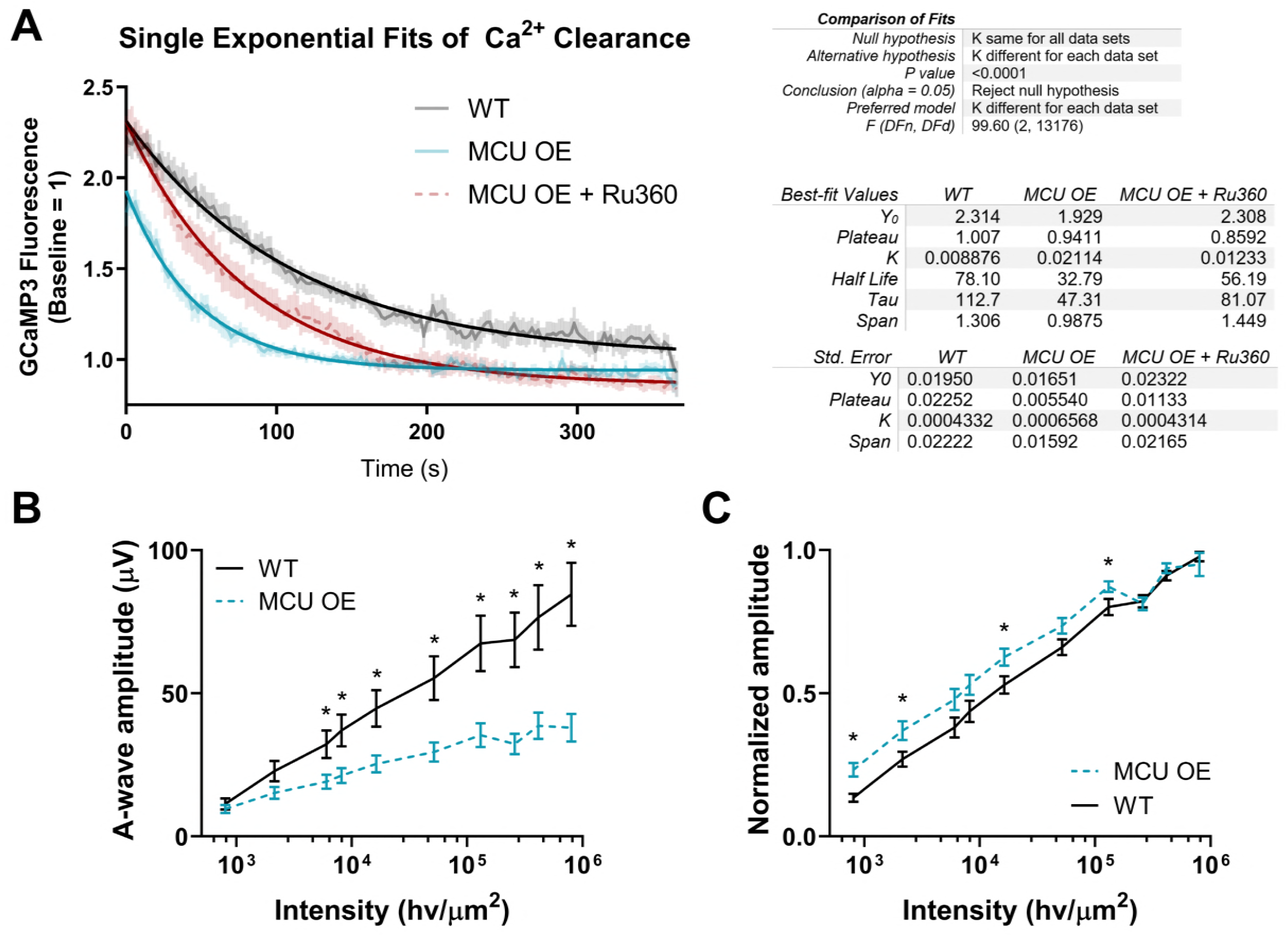
Fitting of Ca^2+^ clearance data and other ERG parameters. A. Truncated Ca^2+^ clearance data used for fitting with a one-phase exponential decay function (least squares fit) in GraphPad Prism 8.0.1. The fitted exponentials are shown by a solid, dark line. No constraints were included for Y_0_ or the plateau, K > 0. K is different for each data set, with p <0.0001. Descriptive statistics of the fit included in table. B. Absolute amplitude of the isolated a-wave response to varying intensity light of WT and MCU OE retinas for experiments shown in Figure 4D. Bars = standard error. *p<0.05 using Welch’s t-test. C. Amplitude of a-wave responses in WT and MCU OE retinas normalized to the maximum response for experiments shown in Figure 4D. Bars = standard error. *p<0.05 using Welch’s t-test.

**Supplemental Figure 5:**
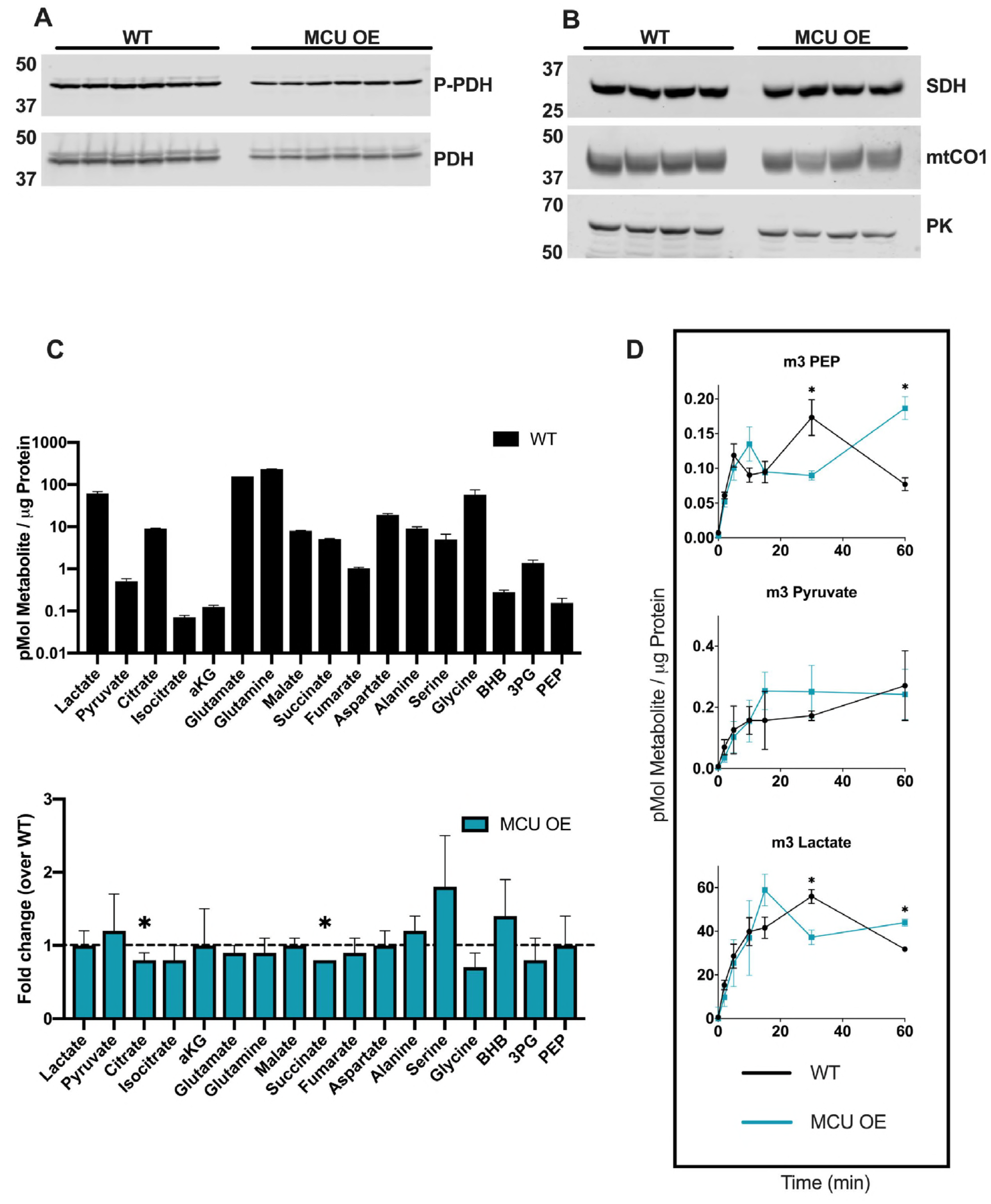
Total metabolites and glycolytic flux in MCU OE retinas. A. Immunoblot showing P-PDH and total PDH expression in WT and MCU OE retinas. n=6 WT and 6 MCU OE retinas from 3 different fish. B. Immunoblot of mtCO1, SDH, and Pyruvate Kinase (PK) in WT and MCU OE retinas. We observed that expression of every protein we probed for (PDH, SDH, mtCO1 and pyruvate kinase) was slightly lower in MCU OE retinas, even when the same amount of protein lysate was loaded. We hypothesize that this is due to the fact that in MCU OE retinas, MCU and RFP comprise a much larger fraction of the total protein, so other proteins appear less abundant when normalizing to total protein. n=4 WT and 4 MCU OE retinas from 4 different fish. C. Total metabolite levels in freshly dissected WT zebrafish retinas and relative levels of these metabolites in MCU OE retinas. (BHB: β-hydroxybutyrate, 3PG: 3-phosphoglycerate, PEP: phosphoenolpyruvate). D. Glycolytic intermediates from WT and MCU OE retinas supplied with ^13^C-glucose. We observed no trends of altered glycolytic flux over time between WT and MCU OE retinas. * p < 0.05 using Welch’s t-test.

**Supplemental Figure 6:**
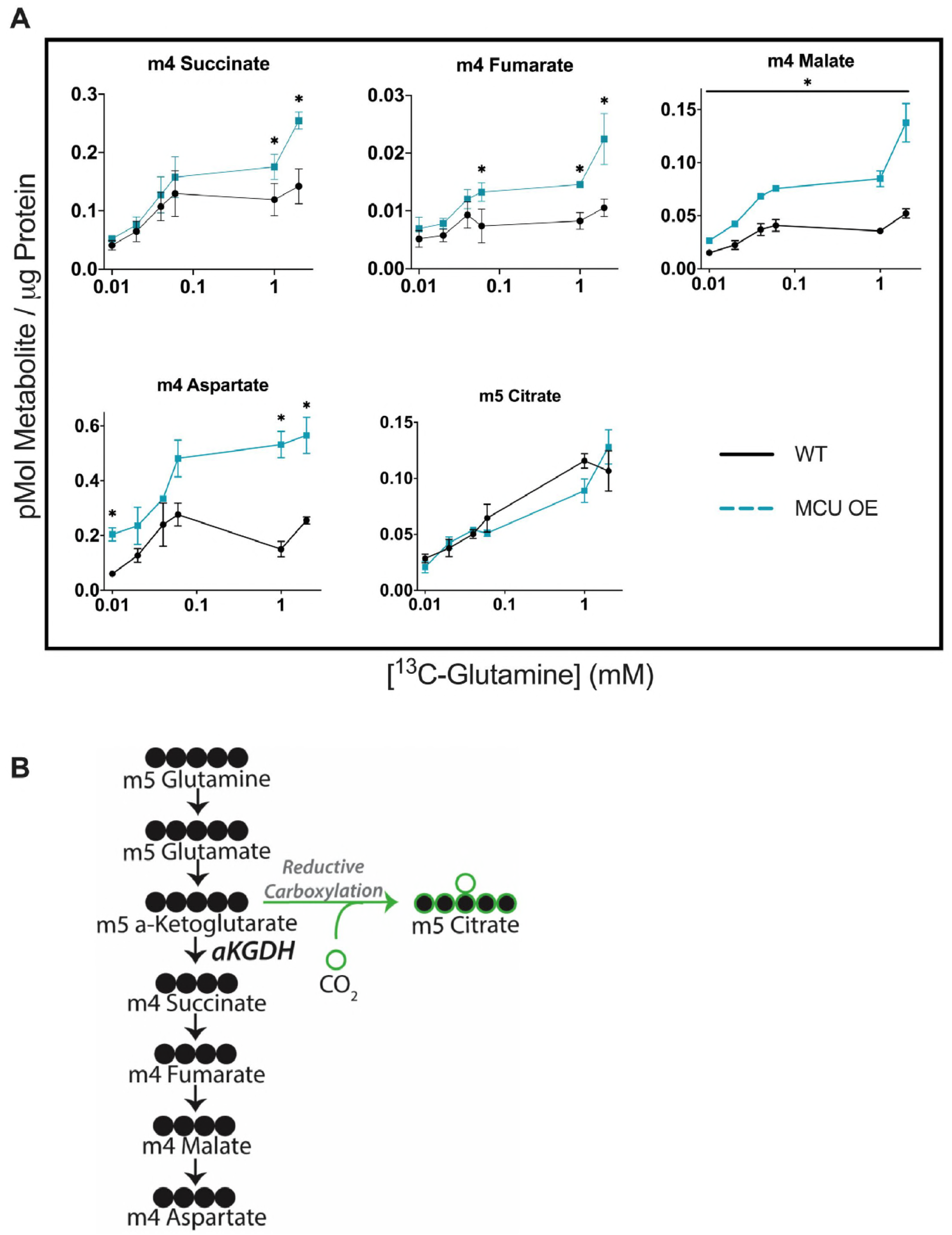
^13^C-Glutamine titration in WT and MCU OE retinas. A. Titration of WT and MCU OE retinas supplied with ^13^C-glutamine (0.1, 0.2, 0.4, 0.6, 1, and 2 mM) for 15 minutes. m5 citrate (produced from reductive carboxylation) is included to show that only metabolites directly downstream of α-KGDH are produced at higher levels in MCU OE cones. Data points represent averages from n=3 retinas from 3 different fish.*p<0.05 using Welch’s t-test. B. Diagram showing how labelled carbons from U-^13^C-glutamine are incorporated into *α*-ketoglutarate and downstream metabolites, including through reductive carboxylation (green outline).

**Supplemental Figure 7.**
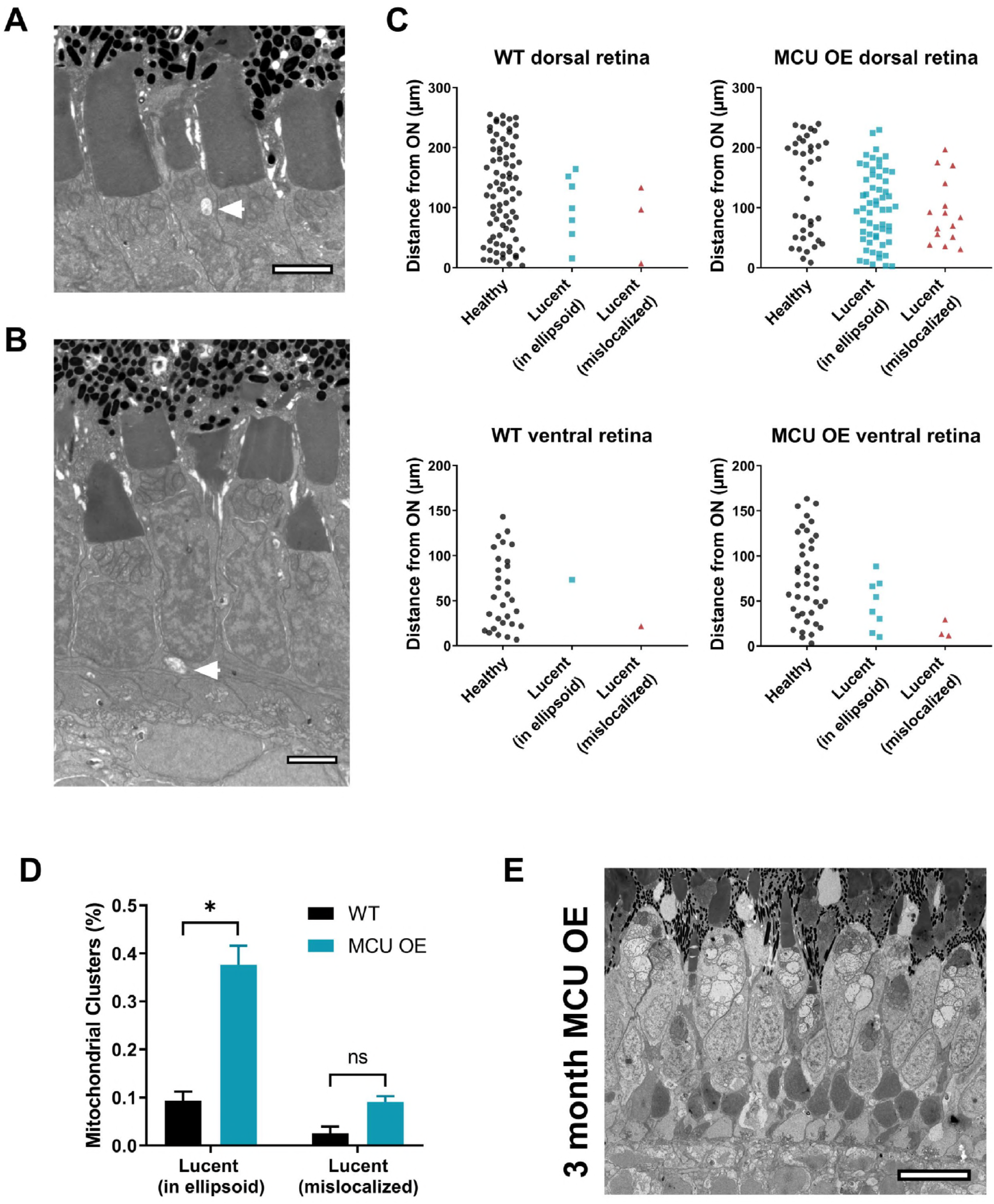
Further characterization of electron lucent mitochondria in WT and MCU OE retinas. A. Electron micrograph from WT fish at 5 days of age. Structures similar to the electron lucent mitochondria in MCU OE fish can be sparsely observed in the ellipsoid WT cones (white arrowhead). These were markedly smaller than structures seen in MCU OE cones. Scale bar = 2 μm. B. Electron micrograph from WT fish at 5 days of age. What appear to be electron lucent, mislocalized mitochondria are sometimes observed in WT fish (white arrowhead). Scale bar = 2 μm. C. Quantification of mitochondrial clusters containing all electron dense mitochondria (healthy) and any electron lucent mitochondria either in the ellipsoid or mislocalized. Data compiled from a single slice at the optic nerve of larval zebrafish at 5 days of age. Clusters were counted and plotted relative to the distance (μm) of the cluster from the optic nerve. Graphs shown are representative plots from dorsal and ventral sides of a single larvae, but this analysis was performed on n=3 larvae for each group for quantification in D. D. Quantification of clusters containing lucent mitochondria as a percentage of total clusters observed (Healthy + Lucent (ellipsoid) + Lucent (mislocalized)) from data shown in C. N=3 larvae for WT and MCU OE. *p<0.05 and ns = not significant using Welch’s t-test. E. Electron micrograph of MCU OE fish at 3 months of age. The electron lucent mitochondrial phenotype is preserved as fish age and is not specific to early development. Scale bar = 10 μm.

**Supplemental Figure 8.**
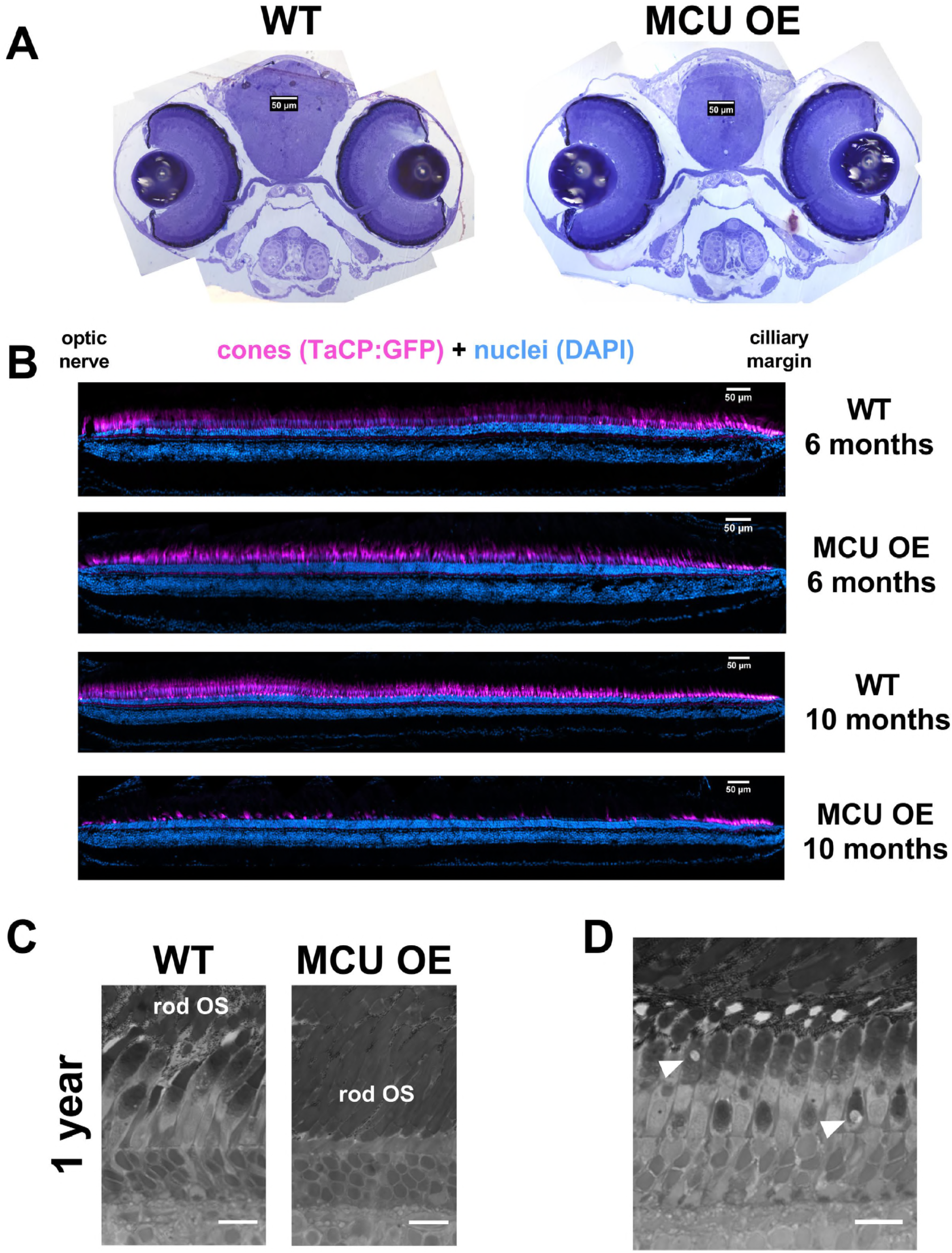
Further characterization of retinal health throughout development in both WT and MCU OE models. A. Larval zebrafish sections at 6 days of age stained with Richardson’s stain. Aside from mitochondrial disturbance in the photoreceptor layer of MCU OE retinas, normal morphology appears to be conserved. Scale bar = 50 μm. B. Representative stitched and straightened images of TαCP:GFP (cones, magenta) retinal sections stained with a nuclear stain (Hoescht, cyan). The double cone nuclei that sit on top of the nuclear layer that contains both UV/blue cones and rods was used for quantification. Scale bar = 50 μm. C. Richardson’s stain of representative 1 year old zebrafish retina from WT and MCU OE fish. In MCU OE models, cones are rarely observed and instead rods have proliferated (rod OS = rod outer segments). Scale bar = 25 μm. D. Richardson’s stain of 1 year old zebrafish retina from WT fish. Occasionally, cone ellipsoid mitochondria clusters in these older fish are observed to have large, lucent mitochondria like younger MCU OE fish (white arrowheads). Scale bar = 25 μm.

